# Amyloid-β increases MBP and MOBP translation in oligodendrocytes through dysregulation of hnRNP A2 dependent RNA dynamics

**DOI:** 10.1101/2024.04.19.590214

**Authors:** Adhara Gaminde-Blasco, Rodrigo Senovilla-Ganzo, Uxue Balantzategi, Fernando García-Moreno, Carlos Matute, Jimena Baleriola, Elena Alberdi

## Abstract

Oligodendrocyte dysfunction, myelin degeneration, and white matter structural alterations are critical events in Alzheimer’s disease (AD) that contribute to cognitive decline. A key hallmark of AD, Aβ oligomers, disrupt oligodendrocyte and myelin homeostasis, but a comprehensive global analysis of the mechanisms involved is lacking. Here, transcriptomic profiling of Aβ-exposed oligodendrocytes revealed widespread gene expression changes, particularly affecting pathways related to RNA localisation. Among the genes identified, we focused on *Hnrnpa2/b1*, the gene encoding the hnRNP A2 protein, which is essential for RNA transport and translation of myelin proteins. We confirmed aberrant upregulation of hnRNP A2 in hippocampal oligodendrocytes from post-mortem human brains of early-stage AD patients, Aβ-injected mouse hippocampi and Aβ-treated disrupting cells *in vitro*. RIP-seq analysis of the hnRNP A2 interactome revealed attenuated interactions with *Hnrnpk* and *Hnrnpa2/b1*, while interactions with *Mbp* and *Mobp* were enriched, suggesting changes in RNA metabolism of molecules associated with mRNA transport of myelin proteins. Aβ increased the total number and dynamics of mRNA-containing granules, facilitating local translation of the myelin proteins MBP and MOBP and attenuating Ca^2+^ signalling. These findings suggest that Aβ oligomers disrupt RNA metabolism mechanisms crucial for oligodendrocyte myelination through dysregulation of hnRNP A2 and myelin protein levels, potentially affecting oligodendroglia Ca^2+^ homeostasis.

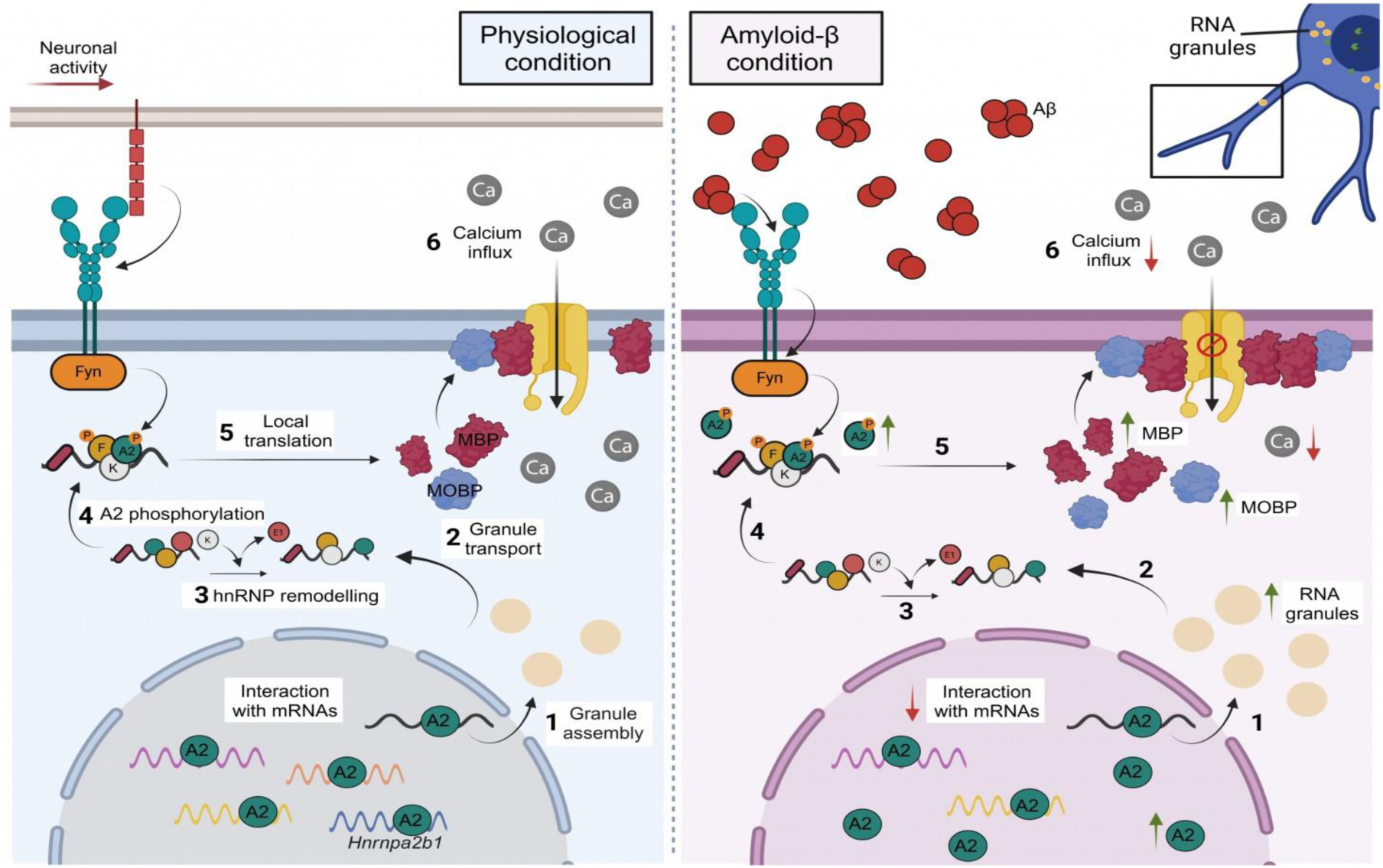
GRAPHICAL ABSTRACT.

## INTRODUCTION

The human brain consists of more than 50% of white matter (WM) (1), which is composed of nerve fibres with varying degrees of myelination that allows transfer of signals across various areas of the central nervous system (CNS). Oligodendrocytes (OLs) are one of the major cell types in the WM. They play a key role by extending membrane processes to wrap around the axons of multiple neurons, creating and maintaining a specialised multilamellar lipid structure known as myelin. Myelin not only prevents ion leakage but also enhances nerve conduction efficiency and speed (2). In addition to insulation, oligodendrocytes provide trophic and metabolic support to axons (3, 4, 5).

The protein content of myelin is dominated by proteolipid protein (PLP) and myelin basic protein (MBP). The formation and maintenance of myelin therefore requires the targeted delivery of large amounts of PLP and MBP to the axon-glia contact site, which is precisely regulated in time and space (6). *Mbp* mRNA is transported from the nucleus within RNA granules to the distal regions of oligodendrocytic processes and to the myelin membrane where it is locally translated preventing ectopic expression within the cell (7). In oligodendrocytes, this process is governed by the heterogeneous nuclear ribonucleoprotein A2/B1 (hnRNP A2 to simplify) with the help of other hnRNPs (8). HnRNP A2 binds to a cis-acting element in the *Mbp* mRNA 3′ untranslated region (UTR) termed the A2 response element (A2RE) (9, 10). Beyond its role in *Mbp* transport and translation, hnRNP A2 is one of the most abundant RNA binding proteins (RBPs) and a key regulator of RNA metabolism, controlling multiple transcripts simultaneously by recognising specific motifs (11).

RBPs are the major regulators of gene expression at the mRNA level, including transcription, processing, transport, and degradation (12). In the nucleus, they regulate mRNA maturation, including RNA helicase activity, RNA polymerase elongation and splicing of multi-exon genes resulting in different protein isoforms from a single gene. They also regulate the amount and timing of mRNA exported to the cytoplasm. In the cytoplasm, RBPs regulate mRNA stability and translation into the correct cytoplasmic localisation (13). The diverse roles of these nucleo-cytoplasmic RBPs suggest that they may act as coordinators of mRNA metabolism.

Recently, a new focus is placed on the role of RNA processing and its impact on neurodegenerative diseases like Alzheimeŕs disease (AD) (14). One of the hallmarks of AD is the presence of extracellular aggregates of amyloid beta peptide (Aβ), and Aβ oligomers-induced changes in oligodendrocytes and myelin (15, 16). Furthermore, myelin and oligodendrocyte abnormalities are associated with increased Aβ peptide levels in AD mouse models (17) and in the brains of AD patients (18, 19). Recently, OLs have been identified as a source of Aβ *in vivo* and thus, active contributors of the disease (20). Moreover, in AD, OLs alterations precede neuronal impairment, and the loss of OLs may lead to cognitive deficits (21, 22, 23). However, the mechanisms by which OLs become dysfunctional and lead to AD pathology remain unknown. *In vitro* studies have shown that Aβ becomes toxic when it oligomerises (24) and can directly alter myelination (25) and damage oligodendrocytes and myelin (26). Aβ oligomers change the MBP expression levels via integrin β1 and Fyn kinase signalling (25), but the functional consequences of MBP overexpression in oligodendrocytes are not yet understood.

Here, we found alterations in RNA metabolism and translation in oligodendrocytes exposed to Aβ oligomers. We reported increased levels of hnRNP A2 in hippocampi of early-stage AD patients as well as in Aβ-injected mice. In addition, we observed that Aβ oligomers increased hnRNP A2 expression and modified associated RNAs directly bound to ribonucleoproteins. This increase in hnRNP A2 coincided with enrichment of *Mbp* and *Mobp* transcripts and an increase in mRNA transport granules that facilitate local translation. Finally, we reported that MBP upregulation reduced Ca^2+^ influx in oligodendrocytes upon plasma membrane depolarisation.

## MATERIAL AND METHODS

### Oligodendrocyte culture and transfections

Highly enriched OPCs were prepared from mixed glial cultures obtained from newborn (P0–P2) Sprague–Dawley rat forebrain cortices as previously described (27) with minor modifications (28). Briefly, forebrains were removed from the skulls and the cortices were enzymatically and mechanically digested. Then, tissue was plated in Iscove’s Modified Dulbecco’s Medium supplemented with 10% fetal bovine serum. The mixed glial cells were grown in poly-D-lysine treated T75 flasks until they were confluent (8–10 days). Microglia were separated from the cultures by shaking the flasks on a rotary shaker. OPCs were isolated by additional shaking for 18 h (200 rpm, 37°C, 5% CO_2_). OPCs were seeded on to poly-D-lysine-coated coverslips and were maintained at 37°C and 5% CO_2_ for 3 days in a chemically defined maturation SATO medium.

In transfection assays, primary oligodendroglial cells were nucleofected with 20 µM siRNA pools (non-targeting siRNA, Dharmacon #D-001810-10-05; *Mbp*-targeting siRNA, Dharmacon #L-089810-02-0005) using Amaxa^TM^ Basic Nucleofector^TM^ kit (#VPI-1006, Lonza) for Primary Mammalian Glial cells following the manufactureŕs instructions.

### Preparation of amyloid β-peptides

Oligomeric amyloid-β (Aβ_1–42_) was prepared as reported previously (29). Briefly, Aβ_1–42_ (Bachem, Germany) was initially dissolved in hexafluoroisopropanol (Sigma) to a concentration of 1 mM. For the aggregation protocol, the peptide was resuspended in dry dimethylsulfoxide (DMSO; Sigma) (Aβ_1–42_ 5 mM). Hams F-12 (PromoCell) was added to adjust the final peptide concentration to 100 μM to obtain oligomers (4 °C for 24 h). In all experiments, Aβ refers to oligomeric amyloid-β unless otherwise stated. As controls, cells were treated with the vehicle (DMSO+Hams F-12).

### Paraffin-embedded human sections

Paraffin-embedded human sections were deparaffinised and rehydrated by immersing in xylene followed by incubations with alcohol solutions (100%, 96% and 75% diluted in H_2_O) in TBS for 10 min each. Samples were then boiled in antigen retriever R-Universal Buffer (Aptum) for 20 min. After retrieval, samples were washed in TBS 3 times and blocked in 4% BSA in TBS for 1 h at room temperature (RT) and incubated overnight (O/N) with the following primary antibodies in blocking solution, hnRNP A2 (sc-374053, 1:1000, Santa Cruz) and Olig2 (MABN50, 1:500, Millipore). Then, samples were washed in TBS twice and incubated with fluorochrome-conjugated secondary antibodies and DAPI (4 µg/ml, Sigma-Aldrich) for 1 h at RT. Samples were washed in TBS and treated with Autofluorescence Eliminator Reagent according to the manufacturer’s instructions (Millipore) to reduce lipofuscin-like autofluorescence. Finally, sections were washed and mounted with Fluoromont-G mounting medium (SouthermBiotech).

### Western blot

Oligodendrocytes were exposed to Aβ peptides as indicated. Cells were scraped in sodium dodecyl sulfate (SDS) sample buffer on ice to enhance the lysis process and avoid protein degradation. Samples were boiled at 95 °C for 8 min, size-separated in 4-20% polyacrylamide-SDS Criterion TGX Precast gels (Bio-Rad) and transferred to Trans-Blot Turbo Midi Nitrocellulose Transfer Packs (Bio-Rad). Membranes were blocked in 5% BSA in Tris-buffered saline/0.05% Tween-20 and proteins were detected by specific primary antibodies against MBP (#SMI 99, 1:1000; Biolegend), MOBP (bs-11184R, 1:1000, Bioss), hnRNP A2/B1 (sc-374053, 1:1000, Santa Cruz), MAG (SC-166849, 1:500, Santa Cruz), MOG (MAB5680, 1:1000, Millipore), CNPase (#C5922, 1:500; Sigma Aldrich), hnRNP F (sc-32310, 1:1000, Santa Cruz), hnRNP K (RN019P, 1:1000, MBL), hnRNP E1 (RN024P, 1:1000, MBL) and β-actin (#A20066, 1:5000; Sigma-Aldrich). Membranes were incubated with horseradish peroxidase-conjugated secondary antibodies (1:5000) or fluorescence secondary antibodies (1:5000). The protein bands were detected with a ChemiDoc XRS Imaging System (Bio-Rad), and quantified by using Image Lab 6.0.1 (Bio-Rad) software.

### Immunochemistry

Cells were fixed in 4% paraformaldehyde, 4% sucrose for 15 min, washed with PBS and blocked in 4% normal goat serum, 0.1% Triton X-100 in PBS (blocking buffer) for 1 h and incubated O/N at 4°C with primary antibodies against MBP (#SMI 99, 1:1000; Biolegend), hnRNP A2/B1 (sc-374053, 1:500, Santa Cruz), hnRNP F (sc-32310, 1:500, Santa Cruz), hnRNP K (RN019P, 1:500, MBL) and hnRNP E1 (RN024P, 1:500, MBL). Samples were washed in PBS and incubated with fluorochrome-conjugated secondary antibodies (1:500) in blocking buffer for 1 h at RT. Cell nuclei were detected by incubation with DAPI (4 μg/ml, Sigma-Aldrich) and mounted with Fluoromount-G (SouthermBiotech).

### Immunoprecipitation

Immunoprecipitation assays were performed to determine changes in hnRNP A2 phosphorylation levels. Briefly, 1 µg of agarose conjugated mouse anti-pTyr (PY99, sc-7020 AC, Santa Cruz) and mouse anti-IgG (sc-2342 AC, Santa Cruz) were used. Cells were washed with cold PBS and scraped with RIPA lysis buffer (ThermoFisher Scientific) supplemented with protease and phosphate inhibitor cocktail (ThermoFisher Scientific).

Samples were placed on ice for 10 min and centrifuged 5 min at 12,000 x g. The supernatant was added (1/10 was saved for input) to the agarose conjugated with anti-pTyr or anti-IgG and incubated for 2 h at 4°C to avoid unspecific interactions. The lysate-antibody-beads complex was centrifuged at 2,000 x g for 2 min and washed three times with RIPA lysis buffer and once by PBS followed by centrifugation to obtain the immunocomplex. Finally, protein elution was carried out in 2x sample buffer after boiling the samples at 95°C for 5 min and centrifuged at 12,000 x g for 1 min. Proteins were analysed by western blot.

### RNA-immunoprecipitation (RIP)

RIP was performed as previously described with some modifications (30). In brief, cells were scrapped in polysome lysis buffer (100 mM KCl, 5 mM MgCl_2_, 10 mM HEPES pH 7.0, 0.5% NP-40, 1 mM DTT, 100 units/ml RNase OUT, 1X Protease Inhibitor Cocktail) and incubated with a suspension of 1 µg agarose conjugated mouse anti-hnRNPA2 antibody (sc-374053 AC, Santa Cruz) and mouse anti-IgG (sc-2342 AC, Santa Cruz). RNA was isolated following a TRIzol RNA isolation protocol (ThermoFisher Scientific). An equal volume of extracted RNA from each sample was then used for cDNA synthesis and analysed by quantitative PCR. Data were normalised to a normalisation factor obtained in geNorm Software through the analysis of the expression of two housekeeping genes (31). *Mbp, Mobp* and *Tau* mRNA enrichment was examined by relativising the RIP fraction values to the normalised input values.

For, the hnRNP A2 interactome analysis, total RNA from three biological replicates of hnRNP A2 and IgG IPs, as well as three replicates of input mRNA were evaluated using Agilent RNA 6000 Pico Chips (Agilent Technologies, Cat. #5067-1513). Sequencing libraries were prepared using “SMARTer Stranded Total RNA-seq Kit v2 – Pico Input Mammalian” kit (Takara Bio USA, Cat. # 634411), following “SMARTer Stranded Total RNA-seq Kit v2 – Pico Input Mammalian User Manual (Rev. 050619)”.

### Quantitative RT-PCR (RT-qPCR)

Total RNA was extracted from cultured oligodendrocytes using RNA Mini Kit (Quiagen) or TRIzol (ThermoFisher Scientific) according to manufactureŕs instructions. Strand complementary DNA synthesis was carried out with reverse transcriptase Superscript TMIII (Invitrogen) using random primers. Specific primers for *Mbp, Mog, Mag, Hnrnpa2b1, Hnrnpf, Hnrnpe1* and *Hnrnpk* were obtained from Qiagen. They were newly designed for *Mobp-81a* (5’-CGCTCTCCAAGAACCAGAAG-3’ and 5’-GCTTGGAGTTGAGGAAGGTG-3’) and *Cnp* (5’-CGCCCACTCATCATGAGCAC-3’ and 5’-CCTGAGGATGACATTTTTCTGAAGA-3’). RT-qPCR reactions were carried out with 25 ng of reverse transcribed RNA and 300 nM of primers diluted in Sybr Green Master Mix reagent (Bio-Rad). PCR product specificity was checked by melting curves. Data were normalised to a normalisation factor obtained in geNorm Software through the analysis of the expression of three housekeeping genes.

### RNA-seq

RNA quantity and quality was assessed using Qubit RNA assay kit (Invitrogen) and Agilent 2100 Bioanalyzer (Agilent RNA 6000 Pico Chips). Sequencing libraries were prepared using “SMARTer Stranded Total RNA-seq Kit v2 – Pico Input Mammalian” kit (Takara Bio USA, Cat. #634411). The protocol was started with 4-10 ng of total RNA. Paired-end readings were generated from all libraries using HiSeq200 sequencer following Illumina protocols. The base calls or BCL were converted into FASTQ files by utilizing Illumina Inc.’s package bcl2fastq and quality control analysis was performed (32) Alignment was carried out with STAR v2.7.1 (33) against Ensembl genome of *Rattus norvegicus* (BN7.2.toplevel ang gtf v105) and expression counts were obtained with htseq-count (-sreverse) (34). The count matrix was imported to R v4.2.2, where expression levels were normalised and further analysed with DESeq2 (35). Those genes with < 5 for the effect of the Aβ-treatment analysis and <2 counts for the interactome analysis in more than one sample per group were discarded.

The impact of Aβ in oligodendrocytes whole transcriptome was explored with the following design: ∼ SV1 + SV2 + Pair + Experiment + Condition + Experiment:Condition. SV1 and SV2 factors have computed based on Surrogate Variable Analysis (36) to extract variability due to unknown noisy factors as Aβ oligomerization and others. To explore the effect of Aβ in the whole-cell transcriptome, the following contrast was used: ‘Condition_Ab_vs_C’. Those genes with p-adjusted value < 0.05 were considered differentially expressed genes.

To obtain the A2 interactome, we split our control and Aβ-treated samples in the analysis, but following a shared design model ∼ Pair + Experiment. Similarly to other interactome literature (37, 38, 39, 40), contrast list (‘Experiment_RIP_vs_Input’, ‘Experiment_IgG_vs_Input’) was carried out. Those genes with p-adjusted value < 0.05 and log2FC > 0 were identified as significantly interacting with A2.

For data visualization and functional enrichment: ggplot2 (41), clusterProfiler (42, 43) and ggVennDiagram (44) were used. The code used can be found in https://github.com/rodrisenovilla/Garminde-Blasco2024.

### Puromycylation-proximity ligation assay (Puro-PLA)

To detect newly synthesised proteins, cells were exposed to puromycin (2 µM, Sigma-Aldrich) for 10 min in the absence or presence of the protein synthesis inhibitor anisomycin (40 µM) for 25 min. After incubation, cells were washed with cold PBS with digitonin and fixed in 4% PFA, 4% sucrose in PBS for 15 min. PLA was conducted following Duolink® PLA Protocol (Sigma-Aldrich). Briefly, as soon as cells were permeabilized and blocked, these were incubated at 4°C O/N with rabbit anti-MBP (A1664, 1:500, ABclonal), rabbit anti-MOBP (bs-11184R, 1:500, Bioss) and mouse anti-puromycin (MABE343, 1:500, Millipore). Detection of newly synthesised MBP and MOBP through PLA was performed according to the manufacturer’s recommendations, using rabbit PLA plus and mouse PLA minus probes and red Duolink detection reagents (Sigma-Aldrich). Finally, cells were washed and treated with Alexa Fluor^TM^ 488-conjugated phalloidin (A12379, ThermoFisher) in 1% BSA in order to make the cytoskeleton visible. Coverslips were mounted on glass slides with Duolink® In situ Mounting Media with DAPI. Images were taken with a Zeiss Apotome 2 (Zeiss) epifluorescence microscope using 63X oil-immersion objective. Image analysis was performed using ImageJ/Fiji software. Phalloidin signal was processed with a Gaussian Blur plugin to create a mask. Employing a “Concentric Circles” plugin, concentric circles at 10 µm intervals emerging from the centre of the cell body were generated and mean values of fluorescence intensity of PLA probes were obtained.

### Calcium imaging

Measurement of cytosolic Ca^2+^ was carried out as previously described (45). Briefly, oligodendrocytes were loaded with 1 μM Fluo-4 AM (Invitrogen) in culture medium for 30 min at 37°C and then washed. Oligodendrocytes were exposed to 25 mM KCl and images were acquired through a 40X oil objective with an inverted Zeiss LSM800 confocal microscope (Zeiss) at an acquisition rate of 1 frame/15 s for 5 min. For data analysis, a homogeneous population of 10-20 cells was selected in the field of view and cell somata were selected as ROIs. Background values were always subtracted and data are expressed as F/F0 ± S.E.M (%) in which F represents the fluorescence value for a given time point and F0 represents the mean of the basal fluorescence level.

### Statistical analysis

All data were expressed as mean ± S.E.M. Statistical analyses were performed using absolute values. GraphPad Prism 8.2.1 software was used applying one-way or two-way analysis of variance (ANOVA) followed by Dunnett’s, Sidak’s and Tukeýs post hoc tests for multiple comparisons and two-tailed, unpaired or paired Student’s t test for comparison of two experimental groups. Results from independent patients, animals or separate experiments were treated as biological replicates (n≥3). Statistical analysis of Aβ treatment and interactome were conveniently explained in the RNA-seq section.

## RESULTS

### 1. Transcriptomic profiling of Aβ-treated oligodendrocytes revealed alterations in RNA localisation and ribonucleoproteins

To test the impact of Aβ oligomers on the oligodendrocyte transcriptome, we performed bulk RNA-sequencing (RNA-seq) on both control and Aβ-treated (1 µM, 24 h, n=3) primary cultured cells. We identified 3,204 differentially expressed genes (DEGs) using DeSeq2 (35) (Padj<0.05; Figure 1A). Among these, 2,002 genes were upregulated, while 1,202 were downregulated (Figure 1A, Supplementary Table 1).

**Figure 1.**
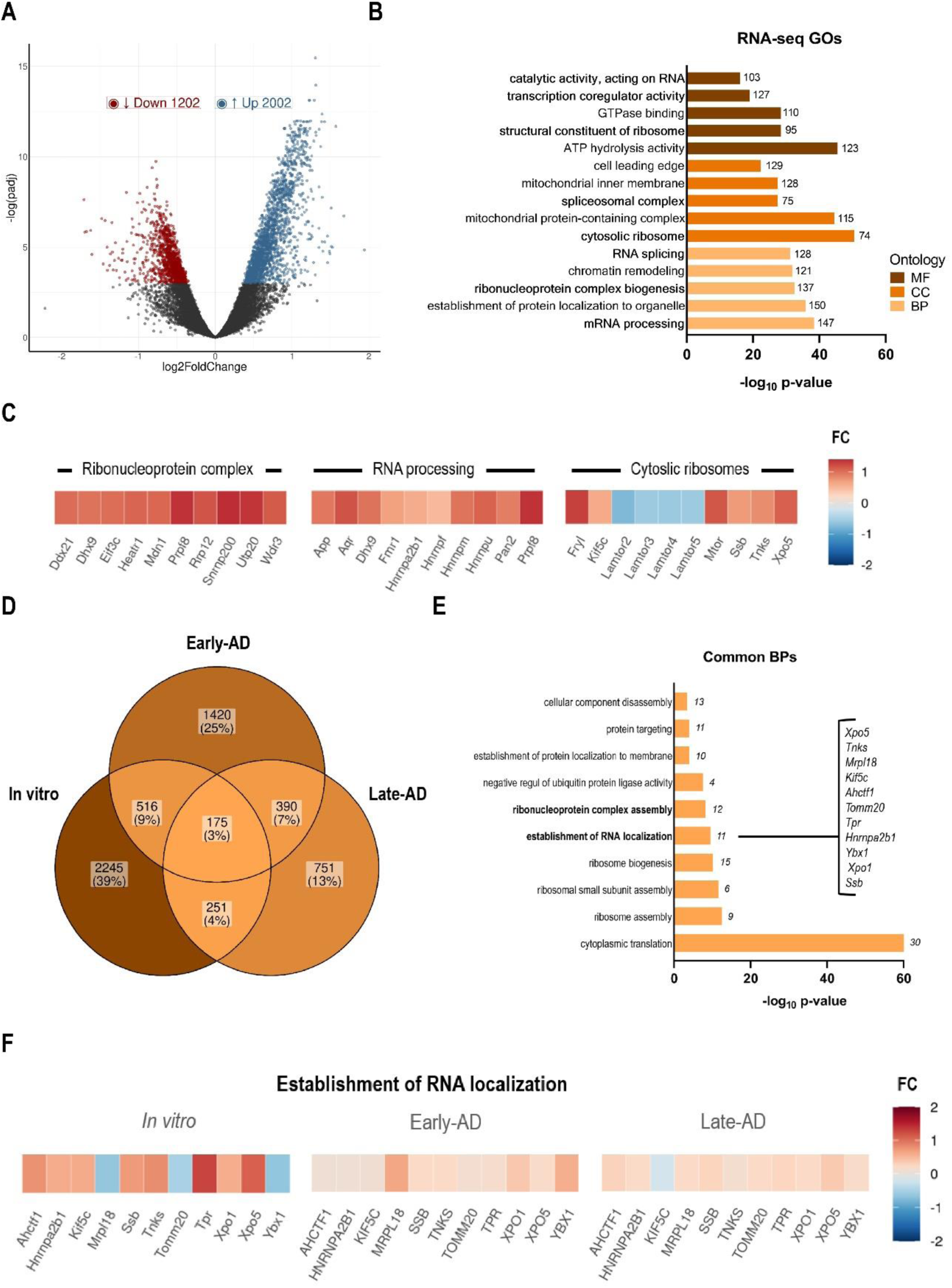
Identification of deregulated genes in AD. (A) Volcano plot of differentially expressed genes (DEGs). Analysed genes are scattered by their expression levels (axis X, log10 mean of normalised counts) and fold change (FC) (axis Y, log2FC). Differentially expressed genes (Padj<0, 05) are coloured based on their FC (red <0; blue >0). (B) Gene ontology enrichment analysis of biological process (BP), cellular component (CC) and molecular function (MF) of DEGs and number of genes associated in each GO term. (C) Selected GO terms related to ribonucleoproteins, RNA processing and myelination. Differentially expressed genes (Padj<0. 05) are coloured based on their FC (red <0; blue >0). (D) Venn diagram depicting the overlap of DEGs *in vitro* vs human AD patients. Percentage and numbers indicate the genes shared among the datasets. (E) Gene ontology enrichment analysis of the biological processes shared in Aβ and AD condition. (F) Selected GO terms related to the establishment of RNA localisation are shown. DEGs (Padj<0. 05) are coloured based on their FC (red <0; blue >0).

To broadly characterise the potential transcriptomic effects of Aβ on molecular processes, we performed Gene Ontology (GO) term enrichment analyses using the Cluster Profiler software (42, 43). Notably, we observed a pronounced enrichment of genes associated with RNA processing (e.g. splicing), ribonucleoprotein complex biogenesis or cytosolic ribosomes (Figure 1B and Supplementary Figure S1 A and B), suggesting a significant impact of Aβ on mechanisms related to RNA metabolism and translation (Figure 1C).

Next, we asked whether the gene signatures associated with Aβ identified *in vitro,* were altered in oligodendrocytes derived from AD patients. We curated published datasets to obtain gene signatures of oligodendrocyte-lineage cells derived from early– and late-stage human AD patients (46). We then analysed the overlap between DEGs obtained from the mentioned datasets and those observed in our *in vitro* model. We identified 175 DEGs common to the three conditions (Figure 1D). Importantly, most of them were primarily linked to cytoplasmic translation, ribosome formation, ribonucleoprotein complex assembly or RNA localisation (Figure 1D and Supplementary Figures S1C and D), confirming the pathophysiological relevance of RNA metabolism and translation in AD.

Many genetic neurological disorders, including AD, are characterised by mutations in RBPs (47, 48), and defects in RNA metabolism (including splicing, localisation or degradation) and translation are common to all of them (14, 49). Such defects have been mainly studied in neurons but given the prominence of DEGs related to the aforementioned mechanisms in oligodendrocytes, we explored the effect of Aβ on RBP levels in cultured oligodendrocytes. Thus, we focused on the 11 RNAs involved in RNA metabolism shared across all three conditions (Figure 1F). Interestingly, among all RNAs, we identified hnRNP A2, one of the most abundant and essential RNA-binding proteins in the central nervous system (CNS), which is known to regulate splicing events in Alzheimer’s disease (50). Furthermore, hnRNP A2 is involved in the regulation of RNA transport and translation of two crucial myelin proteins, MBP and MOBP (8). Based on the literature and our own results, we further focused on studying the regulation of nuclear and cytoplasmic hnRNP A2 in oligodendrocytes in AD.

### 2. hnRNP A2 levels are increased in hippocampal oligodendrocytes from early-stage AD patients

Previous studies have shown reduced expression of hnRNP A2 in neurons of the entorhinal cortex of AD patients (50) but increased expression in the hippocampus (51). However, modifications in hnRNP A2 expression in oligodendrocytes in the context of AD are unknown. Therefore, human hippocampal tissues classified according to Braak stages (II, III, IV, VI) (14) (Supplementary Table 2) were immunostained for hnRNP A2 and Olig2, a specific marker for oligodendrocyte lineage cells. Oligodendrocytes exhibited increased levels of nuclear hnRNP A2 levels in Braak II and III stages compared to controls (control 145.9 ± 5.13, n=3, 120 cells; Braak II 168.5 ± 3.62, n=3, 126 cells; and Braak III 168.5 ± 3.62, n=3, 106 cells; arbitrary units, a.u.). In contrast, no significant differences were observed at later stages (Braak IV 135.3 ± 3.66, n=3, 81 cells and Braak V 152.9 ± 4.38, n=3, 909 cells; a.u.) (Figure 2A and B). These changes suggest a critical temporal dynamics of hnRNP A2 expression throughout the course of the disease in oligodendrocytes.

**Figure 2.**
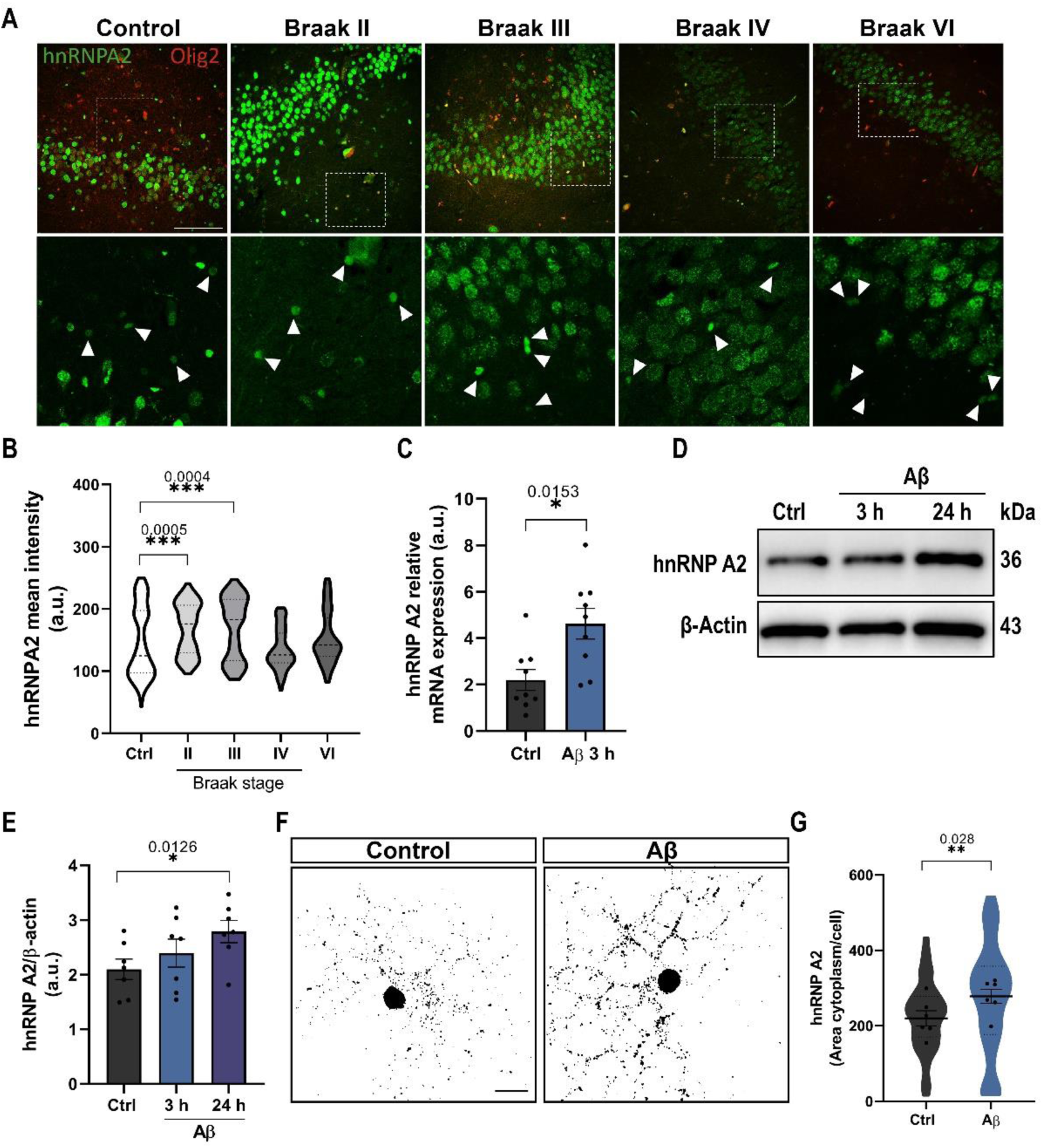
hnRNP A2 expression and localisation in oligodendroglial cells. (A, B) Representative confocal images show the Olig2 (red) and hnRNP A2 (green) expression in the hippocampus from controls and AD patients at Braak II-VI stages. Quantification of hnRNP A2 intensity signal in Olig2+ cells in the human hippocampus. Scale bar 100 µm. (C) RT-qPCR analysis of *hnRNP A2* mRNA expression in Aβ-treated and control cells. (D, E) hnRNP A2 expression and relative quantification in oligodendrocyte cell extracts normalised to β-actin. (F, G) Representative binary micrographs of hnRNP A2 in control and Aβ-treated oligodendrocytes. Histogram depicts changes in the area occupied by hnRNP A2. Data indicate means ± S.E.M and dots represent independent experiments. Statistical significance (*p<0.05, **p<0.01, ***p<0.01) was drawn by two-tailed paired Student t-test (C, G) and two-tailed ordinary one-way ANOVA followed by Dunnett’s post-hoc test (B, E).

### 3. Aβ oligomers increase oligodendrocyte hnRNP A2 expression and subcellular localisation

hnRNP A2 shuttles between the nucleus and the cytoplasm, where it participates in the translation of mRNAs containing the specific cis-acting element, A2RE (9). A more detailed study of Aβ-induced effects on hnRNP A2 was performed in primary oligodendrocyte cultures. We first observed a significant increase in the mRNA levels of hnRNP A2 after Aβ treatment (1 µM, 3 h), as determined by RT-qPCR (control 2.06 ± 0.75 vs. Aβ 3.45 ± 0.57, a.u., n= 9) (Figure 2C). Western blot analysis confirmed a 44.5% ± 12.45 increase in hnRNP A2 protein levels after incubation with Aβ 1 µM for 24 h compared to controls (144.5% ± 12.45 vs. 100% of control, n=7) (Figure 2D and E). On the other hand, immunofluorescence analysis revealed that hnRNP A2 was abundantly present in the soma and processes of the oligodendroglial cells (Figure 2F). Compared to control cells, Aβ treatment caused a significant increase in the granular pattern area (control 21.16 ± 2.03 vs. Aβ 26.95 ± 2.34, n= 6 and 60 cells) in oligodendroglial peripheral processes (Figure 2F and G). To elucidate the biological effect of Aβ on hnRNP A2 protein expression, we injected the peptide or vehicle into the hippocampus of C57 adult mice. Aβ injection (10 µM) led to an upregulation of hnRNP A2 protein in oligodendrocytes in dentate gyrus (DG), as revealed by fluorescence quantification in immunolabelled oligodendrocytes (control 140,772 ± 9,011 vs. Aβ-injected 171,787 ± 8,360, arbitrary units, a.u.; n= 4) (Supplementary Figure S2A and B). Overall, these findings demonstrate that Aβ triggers hnRNP A2 mRNA and protein overexpression in oligodendrocytes *in vitro* and *in vivo* and promotes peripheral localisation of granules in oligodendrocyte processes.

### 4. Aβ oligomers alter hnRNP A2-associated transcripts in oligodendrocytes

hnRNP A2 is primarily localised to the nucleus where it regulates transcription initiation, promotes alternative splicing and facilitates the translocation of mRNAs from the nucleus to the cytoplasm (52). hnRNP A2 exerts all these functions by binding to specific RNA sequences. These interactions of RBPs with their target RNAs are highly dynamic and allow the cell to respond to different environmental conditions. Despite the importance of this protein in health and disease, little is known about its associated transcriptome. To gain insight in the role of hnRNP A2 in oligodendrocytes, we carried out RNA immunoprecipitation sequencing (RIP-seq) using anti-hnRNP A2 and isotype control antibodies (anti-IgG) in primary cultured oligodendrocytes treated with Aβ (1 µM, 24 h, n=3) (Supplementary Figure S3A). Sequencing revealed 1,103 transcripts associated with hnRNP A2 in control cells and 684 transcripts in Aβ-treated cells (DeSeq2, Padj<0.05; Figure 3A and B, Supplementary Excel). Notably, among these RNAs, 655 genes were common to both conditions and 29 were specific to Aβ-treated cells (Figure 3C). In both conditions, 95% were protein-coding transcripts, while 5% were long non-coding (lnRNAs) and 5% were pseudogenes (Figure 3D). We also compared our data with POSTAR3 database (53) and confirmed that, in control conditions, 862 out of the 1,103 RNA targets found in our RIP-seq experiment are described to interact with the hnRNP A/B family.

**Figure 3.**
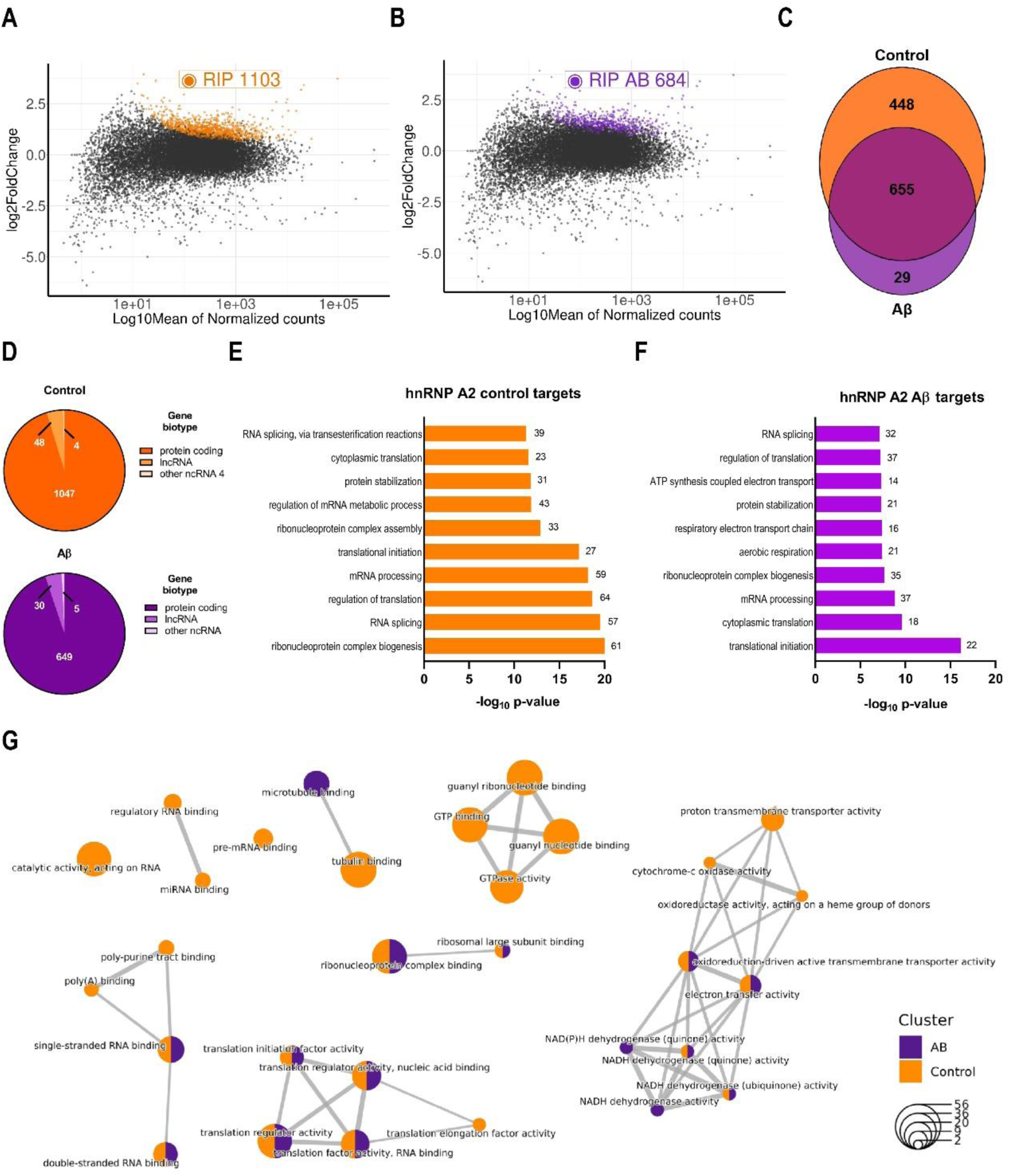
RIP-seq analysis of hnRNP A2-associated RNAs performed in primary cultured oligodendrocytes. (A, B) MA plot of hnRNP A2 RIP-seq data in control and Aβ-treated cells. For each transcript, the average signal (measured as log10 mean of normalised counts) against the RIP-seq log2 Fold Enrichment (RIP versus IgG) was plotted. Significantly enriched targets are highlighted in orange (Ctrl) and purple (Aβ). (C) Venn diagram depicting the overlap between the hnRNP A2 interactome of control (orange) and Aβ-treated (purple) oligodendrocytes. The numbers indicate the genes shared among the datasets. (D) Classification of hnRNP A2-associated RNAs in control (orange) and Aβ-treated (purple) oligodendrocytes. The majority (95%) of identified targets are protein-coding genes, but long noncoding RNAs (lncRNAs) and other ncRNAs were also present. (E) Functional annotation of enriched hnRNP A2 target genes in control (orange) and Aβ-treated cells (purple). The barplot displays the top 10 enriched biological processes. The length of each bar is proportional to the statistical significance of the enrichment. The number of genes associated in each category is displayed beside the corresponding bar. (G) Network representation of the biological processes GO categories is enriched in the hnRNP A2 interactome in control (orange) and Aβ (purple) cells. The size of the node indicates the number of genes in each GO term.

GO Term enrichment (Clusterprofiler) for biological processes revealed that proteins coded by RNAs interacting with hnRNP A2 were significantly associated with ribonucleoprotein complex biogenesis, mRNA metabolism related processes and translational control (Figure 3E). In Aβ-treated cells, most of the GOs were similar, but the abundance of genes was smaller (Figure 3F). Of note, mRNAs encoding proteins involved in metabolic process and GTPases bound to hnRNP A2 in control cells but not in Aβ-treated cells (Figure 3G). Finally, we found hnRNPA2-associated transcripts, which encode components of RBP complexes, including *hnrnpk*, *hnrnpa1*, *hnrnpf* and *hnrnpa2/b1* itself, suggesting an autoregulatory role. Importantly, in Aβ-treated cells, the interaction between hnRNP A2 and *hnrnpk* as well as its own mRNA was lost.

To explore a possible correlation between hnRNP A2-interacting genes altered in Alzheimer’s disease, we conducted a comparative analysis of DEGs in AD (46). Intriguingly, 357 genes (10.19% of 3,503) that interacted with hnRNP A2 were altered in both early and late-stage AD (Supplementary Figure S3B). We then performed GO Term enrichment (Clusterprofiler) for the biological processes which revealed that among the most abundant interacting mRNAs, there were several factors involved in RNA-related processes like ribonucleoprotein complex biogenesis, RNA splicing, RNA processing and mRNA metabolic processes and translational control (Supplementary Figure S4B), consistent with our RIP-Seq results.

### 5. Aβ oligomers modify the composition and dynamics of mRNA-containing granules and activate them by phosphorylation of hnRNP A2

Expression of MBP is a key step during oligodendrocyte maturation and hnRNP A2 is a central component of mRNA-containing cytoplasmic granules, binding directly to the A2RE in the 3′UTR of *Mbp* and *Mobp* mRNAs (9, 54). We previously demonstrated that Aβ oligomers upregulate local translation of *Mbp* through integrin β1 and Fyn kinase signalling (25). This led us to investigate whether the upregulation of hnRNP A2 and its translocation to the cytoplasm could affect the local translation of *Mbp* and *Mobp*.

Since we observed increased hnRNP A2 expression following Aβ treatment, we first explored if Aβ might affect the levels of other hnRNPs. To address this, we analysed both the mRNA and protein levels of various hnRNPs (hnRNP F, hnRNP E1, hnRNP K) typically found in *Mbp* and *Mobp* RNA granules. However, our analysis did not reveal any significant changes in either mRNA or protein levels for any of these hnRNPs (Supplementary Figure S4A-C, n=5). RNA granules are complex, dynamic structures that undergo multiple remodelling steps before mRNA translation can occur. As these granules reach the periphery, hnRNP E1 is replaced by hnRNP K, a crucial step for directing the mRNA to the myelin sheath and initiating translation (Supplementary Figure S5A, n≥4) (55). Given our observations of hnRNP A2 upregulation in the cytoplasm and in processes, we asked whether these granules differ in terms of their dynamics and composition. We quantified the co-localisation of hnRNP A2 and hnRNP F and observed a higher number of granules containing both components in Aβ-treated cells (control 120.77 ± 9.09 vs. Aβ 165.58 ± 10.80, n=5 and 50 cells; area, µm^2^) (Supplementary figure S5C and D), suggesting an increase in granule number. Furthermore, Aβ-treated cells exhibited a significant increase in the total number of hnRNP A2-F-K granules compared to control cells (71.87% ± 2.845 vs. 76.60% ± 2.845 respectively, n=5 and 50 cells) (Supplementary figure S5E).

mRNA granules are highly heterogeneous, which means that some of them can contain mRNAs, while others are empty. Therefore, to determine whether an increased number of granules would also contain more mRNA and thus be more likely to be translated, we investigated whether Aβ alters the interaction between hnRNP A2 and *Mbp* or *Mobp*. To address this, we conducted RNA-immunoprecipitation (RIP) assay using anti-hnRNP A2 and its isotype control antibody (anti-IgG) (Figure 4A). Analysis of mRNA levels via RT-qPCR in the RIP fractions revealed a substantial enrichment of *Mbp* (control 0.21 ± 0.20 vs Aβ 0.50 ± 0.31, n=4) and *Mobp* (control 0.30 ± 0.14 vs Aβ 0.62 ± 0.31, n=4) mRNAs in Aβ-treated compared to control cells, while no significant changes were observed in the input (*Mbp* control 1.74 ± 0.50 vs Aβ 1.52 ± 0.28; *Mobp* control 1.57 ± 0.72 vs Aβ 1.30 ± 0.46 (Figure 4B). Notably, we observed minimal expression of *Mbp* (0.017 ± 0.014) and *Mobp* (0.023 ± 0.012) in the control condition with the anti-IgG. Lastly, to determine if the enrichment was specific to *Mbp* and *Mobp*, we investigated *Tau*, which binds to hnRNP A2 (56). No significant changes were observed in the RIP fractions (control 0.17 ± 0.14, Aβ 0.17 ± 0.13, and IgG 0.0014 ± 0.0006, n=4) nor in the input (control 1.64 ± 0.19 vs Aβ 2.01 ± 0.246) (Figure 4B). This suggests that Aβ promotes mRNA-containing granules in oligodendrocytes.

**Figure 4.**
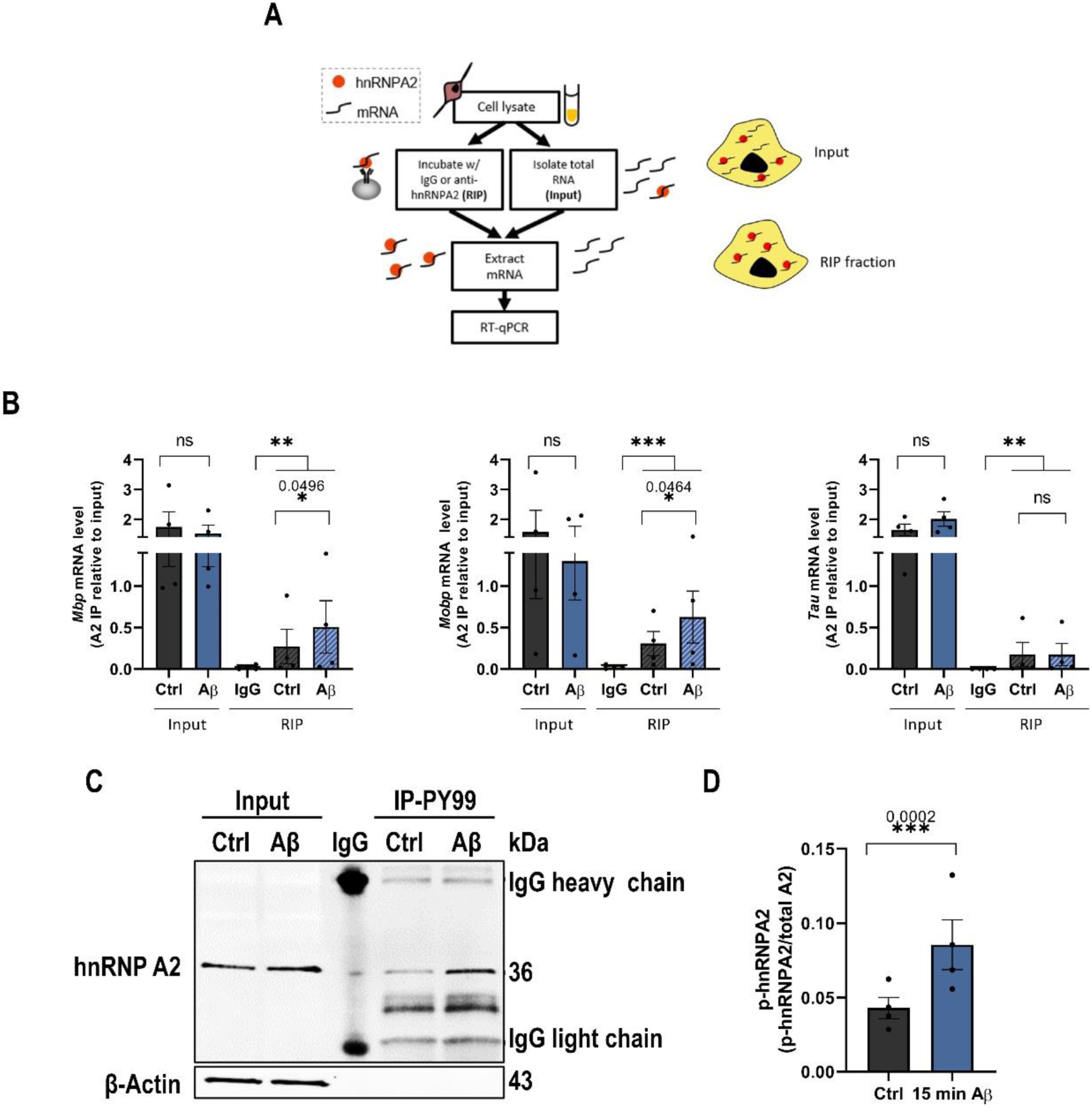
mRNA granule content and activation. (A) HnRNPA2-RIP workflow. Control IgG and anti-hnRNPA2 were used for RIP from oligodendrocyte lysates. (B, C, D) Analysis of *Mbp*, *Mobp* and *Tau* mRNAs levels by RT qPCR (C, D) Western blot of pTYR-IP and IgG to detect hnRNP A2 phosphorylation. (B) Histogram depicting the phosphorylation of hnRNP A2 normalized to total hnRNP A2 from input. Data indicate the means ± S.E.M and dots represent independent experiments. Statistical significance (*p<0.05, **p<0.01, ***p<0.001) was drawn by two-tailed paired Student’s t-test (B, D).

To initiate translation, hnRNPA2 must be disassembled from the mRNA granule through phosphorylation of tyrosine residues by the tyrosine protein kinase Fyn, which requires activation (57). Therefore, to determine if Fyn activation by Aβ increases hnRNP A2 phosphorylation, we conducted an immunoprecipitation assay using an anti-pTyr (PY99) antibody and an isotype control antibody (Figure 4C), followed by a western blot with hnRNP A2 antibody. Data demonstrated that after Aβ treatment (1 µM, 15 min, n=4), the levels of phosphorylated hnRNP A2 were significantly higher than those observed in the control (Figure 4D) (0.042 ± 0.007 vs 0.085 ± 0.016, respectively).

### 6. Aβ oligomers promote local translation of *Mbp* and *Mobp*

To investigate whether the observed alterations in RNA granules and hnRNP A2 phosphorylation promote the translation of *Mbp* and *Mobp* mRNA, we used puromycin together with proximity ligation assay (Puro-PLA) to assess translation. To visualise oligodendrocyte processes, the cytoskeleton was labelled with phalloidin, which binds to F-actin. We quantified the PLA intensity in both primary processes (those emerging from the soma) and total processes to determine if there were differences in localisation (Figure 5A). Our findings revealed that Aβ-treated oligodendrocytes exhibited significantly enhanced local translation of *Mbp*, both in primary processes (control 162.50 ± 63.66 vs Aβ 262.80 ± 76.43, n=4) and total processes (control 54.34 ± 18.56 vs Aβ 89.74 ± 21.96, n=4) (Figure 7B). However, the results for *Mobp* local translation were less definitive: while Aβ induced increased local synthesis of MOBP in primary processes (control 6.43 ± 2.84 vs Aβ 140.35 ± 104.04, n=3), no significant difference was observed in total processes (control 42.60 ± 27.28 vs Aβ 55.55 ± 37.35, n=3) (Figure 5B). To confirm the translation dependency of the observed puncta in puro-PLA images, we included anisomycin as a negative control, administered 20 minutes prior to puromycin to inhibit translation initiation. As depicted in the puro-PLA images, cells preincubated with anisomycin did not exhibit any PLA-positive puncta, confirming that the puncta we observed in puro-PLA were indeed translation dependent.

**Figure 5.**
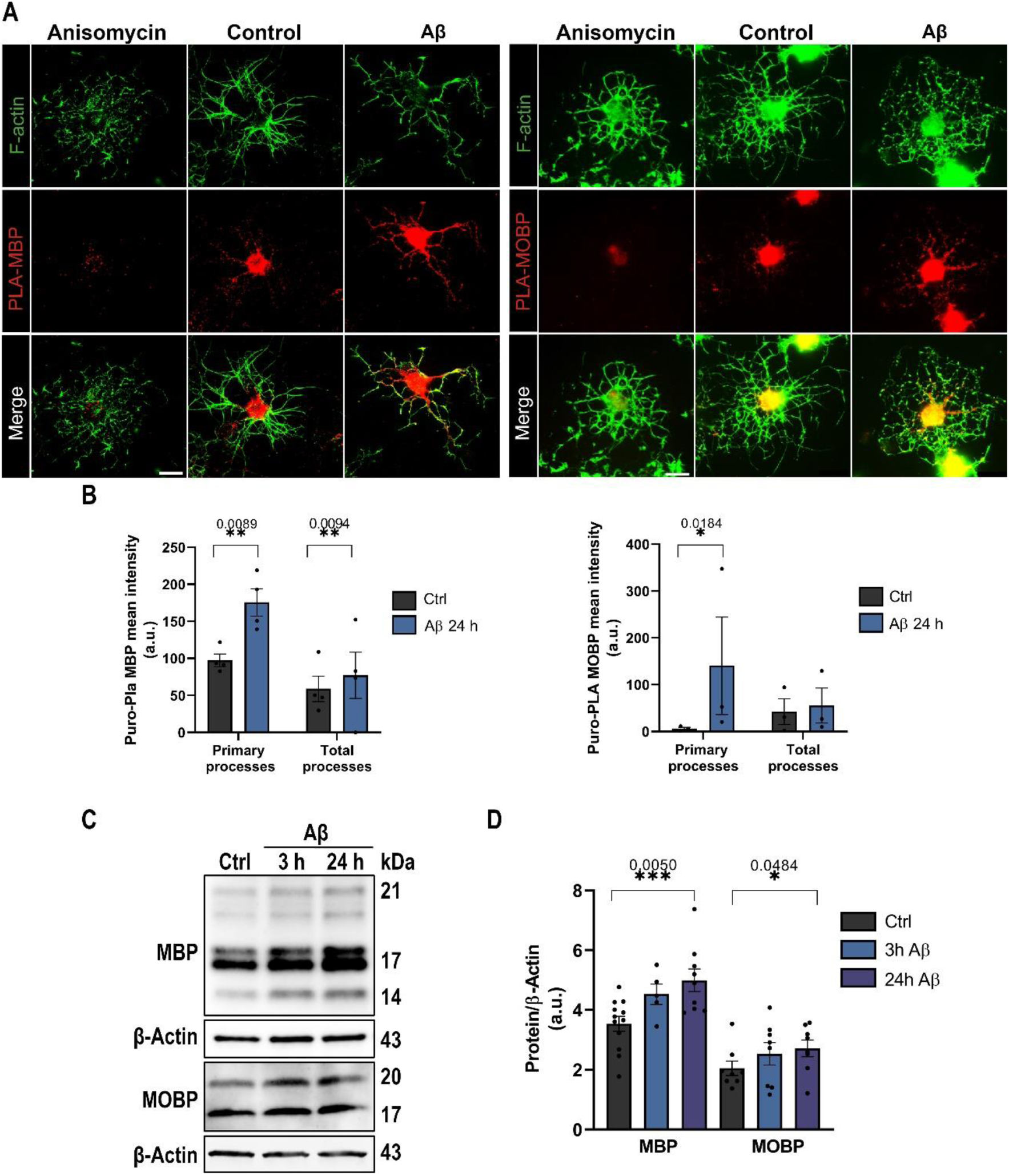
Local translation of Mbp and Mobp. (A) Photographs show MBP and MOBP puro-PLA-positive puncta in the soma, primary and total processes. (B) MBP and MOBP PLA positive puncta was analysed in bins of 10 μm ranging from the soma in primary and total processes. Scale bar, 10 µm. (C, D) MBP and MOBP expression and relative quantification in treated and control oligodendrocyte cell extracts. Data are represented as means ±S.E.M and dots indicate independent experiment. Statistical significance (*p<0.05, **p<0.01, ***p<0.001) was drawn by two-tailed paired Student’s t-test (B) and one-way ANOVA followed by Dunnett’s post-hoc test (D).

We further validated these results through RT-qPCR and western blot analyses, not only for locally translated proteins (MBP and MOBP) but also for other crucial myelin-related proteins such as PLP, CNPase, MOG, and MAG. While we did not observe significant differences in the mRNA levels of *Plp, Cnp, Mog*, or *Mag* (Supplementary Figure S6A, n≥4), we did note that the levels of *Mbp* (1.26 ± 0.21 vs 2.64 ± 0.70 respectively, n=5) and *Mobp* (0.70 ± 0.16 vs 0.84 ± 0.20 respectively, n=8) mRNA increased after 3 hours of Aβ treatment compared to the control (Supplementary Figure S6A). In contrast, analysis of various myelin-related proteins by western blot revealed that MBP (150% ± 13.45, n≥5), MOBP (138.3% ± 17.46, n≥6), and MOG (111.6% ± 3.90, n=6) levels significantly increased after 24 hours of Aβ treatment compared to the control, represented as 100% (Figure 5C and D, Supplementary Figure S6B and C). In summary, these results collectively suggest that Aβ not only increase the quantity of RNA granules but also modifies their composition in favour of molecular components that facilitate mRNA translation.

### 7. MBP overexpression by Aβ oligomers reduce calcium dynamics in oligodendrocytes

MBP is a multifunctional protein that interacts with lipids and a diverse array of proteins (58, 59). Specifically, MBP regulates voltage-gated Ca^2+^ channels (VGCCs) at the plasma membrane reducing the Ca^2+^ influx in the oligodendrocyte cell line N19 as well as in primary cultures of oligodendroglial progenitor cells (60). To obtain a direct correlation between Aβ-induced MBP and Ca^2+^ dysregulation, we knocked down the expression of MBP in primary cultures of oligodendrocytes by using small interfering RNAs (siRNAs) against *Mbp* and a non-targeting siRNA as a negative control. Initially, we validated the reduction in *Mbp* gene expression using western blot analysis of 21, 17 and 18 kDa MBP isoforms (Figure 6A and B). The *Mbp*-targeting siRNA decreased the 21 kDa isoform 38.78% ± 10.14 and the 17 and 18 kDa isoforms 19.49% ± 10.50 compared to the negative control (control expression, 100%, n=6) (Supplementary Figure S7A). Aβ-treated oligodendrocytes exhibited an increase in MBP 21 kDa (121.6% ± 7.78) and 17-18 kDa isoforms (132.1% ± 11.92), an effect that was abolished when MBP was inhibited (Supplementary Figure S7A).

**Figure 6.**
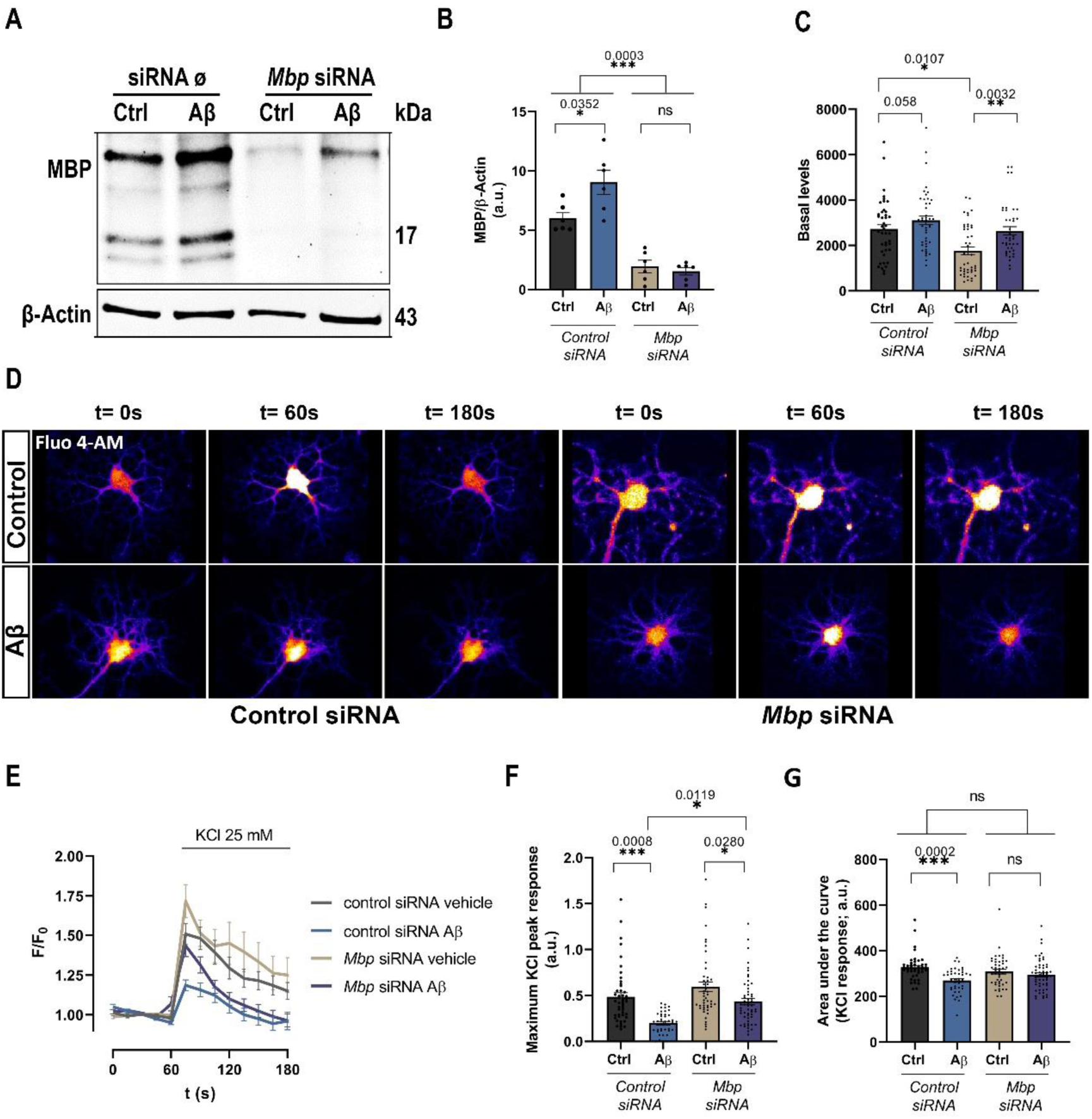
MBP overexpression inhibits KCl-induced Ca^2+^ influx into the cell. (A-B) Western blot and analysis of MBP expression levels following Aβ exposure in oligodendrocytes treated with control siRNA and *Mbp* siRNA are shown (C-E). Cells transfected with either control siRNA or *Mbp* siRNA were loaded with Fluo-4AM and exposed to Aβ for 24 h. Time course of intracellular Ca^2+^ levels were recorded before and after KCl 25 mM stimulus by confocal microscopy. Basal levels are shown in the graph. (F, G) Bar graphs show the maximum peak and the area under the curve (AUC) of KCl response in the different conditions. Data indicate means ± S.E.M and dots represent independent experiments (B, C) and cells (F, G). Statistical significance (*p<0.05, **p<0.01, ****p<0.0001) was drawn by ordinary two-way ANOVA followed by Sidak’s post-hoc test (B, C, F, G).

To investigate whether Aβ modifies Ca^2+^ homeostasis through dysregulation of MBP levels in oligodendrocytes, we performed Ca^2+^ imaging experiments in cells treated with siRNAs targeting and non-targeting *Mbp*. First, we observed reduced levels of resting intracellular Ca^2+^ in *Mbp* knockdown cells compared to controls (Figure 6C), suggesting a role for MBP in Ca^2+^ homeostasis. Next, Ca^2+^ influx by VGCCs after plasma membrane depolarization with KCl was reduced in Aβ-treated compared to control oligodendrocytes (0.448 ± 0.043 vs 0.2091 ± 0.024, n=6 and 47 cells and n=6 and 41 cells), which was partially restored by silencing MBP (0.5967 ± 0.056 vs 0.3966 ± 0.032, n=6 and 47 cells and n=6 and 55 cells) (Figure 6E-G). Moreover, we observed a decrease in the area under the curve (AUC) in Aβ-treated cells (313.7 ± 5.49 vs 272.8 ± 7.49), which was fully recovered when MBP was reduced (300.7 ± 8.37 vs 289.1 ± 8.36) (Figure 6F). These results indicate that MBP overexpression inhibits Ca^2+^ influx through VGCCs, suggesting that Aβ plays a role on Ca^2+^dynamics by modulating MBP levels.

To further investigate whether Aβ could alter Ca^2+^ homeostasis, we employed rat hippocampal organotypic cultures as a more complex cellular system, as they partially maintain tissue architecture, anatomical relationships, and network connections. Oligodendrocytes in these cultures were labelled by transducing organotypic slices with rAAV8-pMBP-GFP, with GFP protein becoming evident as early as 72 hours post-infection. GFP fluorescence was observed in the cytoplasm, nucleus and processes of Olig2+ oligodendrocyte cells, but not in NeuN+ neurons and GFAP+ astrocytes (Supplementary Figure S5B). Subsequently, we treated rat hippocampal slices after 7 days *in vitro* (DIV) with Aβ at 1 µM for 48 hours (n=2 and ≥7 cells) (Supplementary Figure S5C) and recorded intracellular Ca^2+^ levels upon KCl stimulation using puff technique. Our observations clearly indicated that Aβ treatment reduced the maximum peak and AUC (0.090 ± 0.002 vs 0.0336 ± 0.006) (Supplementary Figure S5D and E), suggesting a decrease in Ca^2+^ influx through voltage-gated Ca^2+^ channels. Taken together, these findings confirm that Aβ alters Ca^2+^ influx into cells through mechanisms involving regulation of VGCCs.

## Discussion

Emerging evidence highlights the significant role of white matter degeneration in the pathology of Alzheimer’s disease, although the exact causes remain under investigation. Several studies suggest that abnormalities in myelin and oligodendrocytes in AD are associated with elevated levels of Aβ peptides (18, 19, 61). These changes occur before neuronal damage, and oligodendrocyte loss may contribute to cognitive deficits (21, 22, 23).

Recent reports showed common perturbations in RNA metabolism in several neurodegenerative diseases, standing out the critical importance of ribonucleoprotein homeostasis in brain physiology (65). Using a comprehensive approach including immunohistological analysis of post-mortem human brains, bulk and hnRNP A2-RIP transcriptomics of Aβ-treated oligodendrocytes, and functional analysis to monitor myelin protein translation and intracellular Ca^2+^ levels, we have systematically investigated the impact of hnRNP A2 dysregulation on AD models. Our study elucidates changes in mRNA association and trafficking between the nucleus and cytoplasm, as well as relevant changes in myelin proteins, which in turn influence intracellular Ca^2+^ levels.

Several transcriptomic analyses have been performed in AD patients in recent years (46, 62, 63, 64). In all of them, oligodendrocytes showed alterations in several functions, including myelination, sensing of neuronal activity and immune function (65). Furthermore, the proteomic profile of AD brain networks has revealed a consistent upregulation of oligodendrocyte-enriched modules associated with myelination (66, 67), suggesting an active involvement of oligodendrocytes in the pathology of the disease. Emerging evidence indicates that the signaling of Aβ peptide affects the transcriptional profiles of oligodendrocytes. Spatial transcriptomic analyses (68) demonstrated that oligodendrocytes exhibit distinct transcriptomic responses within plaque environments, suggesting that soluble Aβ peptide alters the oligodendrocyte response during the course of the disease. Consistent with these findings, our research demonstrated that Aβ oligomers induce changes in the oligodendrocyte transcriptome, particularly highlighting genes associated with hnRNPs and RNA metabolism, such as *App*, *Hnrnpa2/b1*, *Hnrnpf*, *Hnrnpm*, and *Hnrnpu*. In agreement with these results, proteomic analysis have shown upregulated modules in AD related with RNA binding and splicing proteins (67) with hnRNP A2 emerging as a common hub (66).

RBPs play a crucial role in RNA metabolism processes by forming complexes that regulate pre-mRNA splicing, transcription, and translation. Previous studies have shown that RNA processing is impaired in neurons of AD brains (69, 70). Amongst the most abundant RBPs are the members of the heterogeneous nuclear ribonucleoproteins hnRNP A/B family, namely hnRNP A1 and A2/B1. In our study, we observed increased levels of hnRNP A2 in oligodendrocytes in the hippocampus of early stages of AD and in the Aβ-injected mice. Consistent with our findings, Mizukami et al. reported increased hnRNP A2 levels in the CA2 region of patients with mild cognitive impairment (51) and decreased levels in dementia (50, 71), indicating region– and time-specific responses in AD pathology. Importantly, depletion of hnRNP A2 leads to the production of a more active β-secretase isoform (BACE1), which may contribute to the accumulation of Aβ plaques (72).

In this study, we described for the first time the transcripts associated with hnRNP A2 in oligodendrocytes. In particular, in agreement with data described for hnRNP A1 (73), hnRNP A2 regulated genes associated with RNA biology, involved in RNA metabolism, processing and splicing, biogenesis of ribonucleoprotein complexes and translation regulation. Intriguingly, we observed that Aβ upregulates hnRNP A2 and disrupts its associated transcripts, leading to a weakening of interactions with over 50% of total mRNAs. Reduced RNA binding by hnRNP A2 results in either production of alternatively spliced protein isoforms, or nonsense-mediated decay (NMD) of mRNA with retained intronic sequences (52). Thus, alterations in hnRNP A2 interactions induced by Aβ may exert profound downstream effects on the cellular transcriptome and proteome.

Local protein synthesis within myelin compartments is tightly regulated in space and time in response to neuronal activity and serves as a critical reservoir for myelin renewal. The localisation of *Mbp* and *Mobp* mRNA is largely dependent on their association with hnRNP A2 and the formation of RNA transport granules. We have previously shown that low levels of Aβ promote the synthesis of MBP (25). The tyrosine kinase Fyn is known to stimulate *Mbp* and *Mobp* transcription and local translation to mediate oligodendrocyte maturation (54, 74, 75). In addition, Fyn has been described as a downstream target of Aβ *in vitro* and in animal models (25, 76). Notably, Fyn phosphorylates the granule proteins hnRNP A2 and hnRNP F, thereby facilitating the translation of *Mbp* and *Mobp* (54, 57, 77).

In the present study, we provide the first evidence that Aβ promotes local translation of *Mbp* and *Mobp* by altering the number and dynamics of mRNA granules through regulation of hnRNP A2. We found a significant increase in the number of granules containing hnRNP A2 and F, as well as active granules containing hnRNP K. The exchange of hnRNP E1 for hnRNP K has been suggested to be a prerequisite for targeting mRNA to the myelin sheath or for the recruitment of protein factors required for translation initiation (55). Thus, more granules containing hnRNP K would indicate that more granules are approaching their final destination for translation. We also showed a higher association between hnRNP A2 and *Mbp* and *Mobp* mRNAs, in addition to higher hnRNP A2 phosphorylation.

MBP is one of the most abundant proteins in the myelin sheath and the only structural myelin protein known to be essential for the formation of compact myelin sheaths (78). Numerous studies have indicated that the absence of this protein leads to myelin vesiculation and subsequent breakdown (79). However, there is no study describing the opposite effect. It is therefore plausible that an excess accumulation of this protein might influence myelin compaction, subsequently affecting conduction velocity and the delivery of nutrients to the axon.

Upregulation of MBP inhibits VGCCs, thereby reducing Ca^2+^ influx into oligodendrocytes (60). Ca^2+^ influx through membrane channels is a critical step in signalling pathways involved in the regulation of growth, maturation and functional plasticity. In addition, elevated Ca^2+^ levels stimulate MBP synthesis in oligodendrocytes (80) and are critical for myelin sheath extension (81). Here, we demonstrate that Aβ inhibits the Ca^2+^ influx into the cell through VGCCs, which is partially recovered by siRNA-mediated MBP downregulation. In addition, we have also shown that Aβ inhibits Ca^2+^ entry into oligodendrocytes in organotypic hippocampal cultures. Since Ca^2+^ entry stimulates MBP synthesis and MBP in turn inhibits Ca^2+^ entry, our results may suggest a self-regulatory mechanism in which MBP inhibits VGCCs to regulate its own synthesis. The accumulation of MBP in the membrane would inhibit or reduce the number of VGCCs in the membrane, as previously reported (60), thereby affecting to the calcium entry through these channels. Thus, our results suggest that Aβ oligomers disrupt Ca^2+^ regulation, making oligodendrocytes more susceptible to environmental stimuli that increase intracellular Ca^2+^ levels and affect myelination.

Overall, this study sheds light on novel mechanisms implicating hnRNP A2, MBP and Ca^2+^ dysregulation in oligodendrocyte lineage cells in the pathophysiology of AD. The results suggest that intervention in Aβ-triggered pathways in oligodendrocytes may offer a therapeutic strategy to improve memory function in the early stages of AD.

## Data availability

The data used in this study is available on the GEO repository with the identifier GEO GSE263799.

## Acknowledgements

We thank L. Escobar, A. Pérez-Samartín and Z. Martínez for technical assistance. F. Soria for technical assistance and helpful discussion. We also thank A. Aransay from CIC BioGUNE Genome Analysis Platform for the technical assistance and the preparation and sequencing of the RNA libraries. Graphical abstract was created with BioRender.com.

## Author’s contributions

A G-B designed and performed the experiments, analyzed and interpreted data and wrote the manuscript. R S-G, UB, F G-M and CM performed the experiments, analyzed and interpreted data. JB provided help to design the project, with data interpretation and reviewed the manuscript. EA contributed to the conception and design of the project, analyzed and interpreted data and wrote the paper. All authors have read and approved the final manuscript.

## SUPPLEMENTARY MATERIAL

### SUPPLEMENTARY MATERIAL AND METHODS

#### Intrahippocampal injection in adult mice

10-week-old male mice (C57BL6/J) were subjected to intrahippocampal injections in the right dentate gyrus (DG). For the surgery, animals were anesthetised with 0.3 ml of Avertin, with addition of 0.1 ml if needed. Mice were immobilized with a stereotaxis apparatus and injected with the corresponding preparations: vehicle or 3 µL of Aβ at 10 µM. After injection and before removal, the needle was left on site for 5 min to avoid reflux. Mice were anesthetised with ketamine /xylacine (100/10 mg/kg) and perfused with 30 ml of phosphate buffer followed by 30 ml of 4% PFA in 0.4 M PB. Brains were extracted and postfixed with the same fixative solution for 4 h at RT, placed in 30% sucrose in 0.1 M PBS pH 7.5 at 4°C, and then stored in cryoprotectant solution (30% ethylene glycol, 30% glycerol and 10% 0.4 M PB in dH_2_O) at –20°C.

#### Organotypic hippocampal slice culture

Organotypic hippocampal slice culture were prepared from brain sections of P5-P7 Sprague-Dawley rat pups according to previously described procedures (82). Briefly, after decapitation, brains were extracted and cut into 350 μm coronal slices using a tissue chopper, hippocampi were then separated and meninges were removed. Slices were transferred to 0.22 μm culture membranes inserts (Millipore), each containing two to three slices. Slices were maintained in 6-well plates for nine days in 25% basal medium with Earlés balanced salts (Life technologies), 44% Minimal Essential Medium (Life technologies), 25% inactivated horse serum (Life technologies), 35.2% glucose solution (Panreac), 50 μg/ml fungisone, B-27 supplement (Gibco), Glutamax (Gibco), 1% Penycilin/Streptomycin (Lonza) and AraC 4.4 µM (only at 2 DIV) at 37°C with 5% CO_2_. After two days, organotypic slices were infected with 1 µL of 10^12^ pAAV-8 carrying MBP promoter (AAV-MBP-GFP) bound to GFP. Slices were kept in culture for 9 days before performing the experiments.

#### Calcium imaging in organtopypic hippocampal slice culture

Organotypic slices were loaded with 5 μM Fura 2-AM (Invitrogen) and 0.01% Pluronic TM F-127 (Invitrogen) in culture medium for 30 min at 37°C. Then, cells were washed in HEPES-Hank’s solution (incubation buffer) (HBSS, MgCl_2_ 1 mM, CaCl_2_ 2 mM, Glucose 10 mM, NaHCO_3_ 4 mM and HEPES 20 Mm; pH 7.4) 10 minutes at RT. Experiments were performed in a coverslip chamber continuously perfused (1.5 mL/min) with the incubation buffer. The perfusion chamber was mounted on the stage of a Leica DMLFSA epifluorescence microscope (Leica) equipped with a Polychrome V monochromator (Till Photonics). Organotyic cultures were exposed to a local application of 50 mM KCl through a microinjector (Eppendorf) with an applied pressure of 160 hPa (duration of the pulse 10 s). Images were acquired through a 40X water objective at an acquisition rate of 1 frame/s during 2 min with Aquacosmos imaging software (Hamamatsu). Intracellular calcium levels were estimated by the 340/380 ratio method.

## SUPPLEMENTARY FIGURES

**Figure S1.**
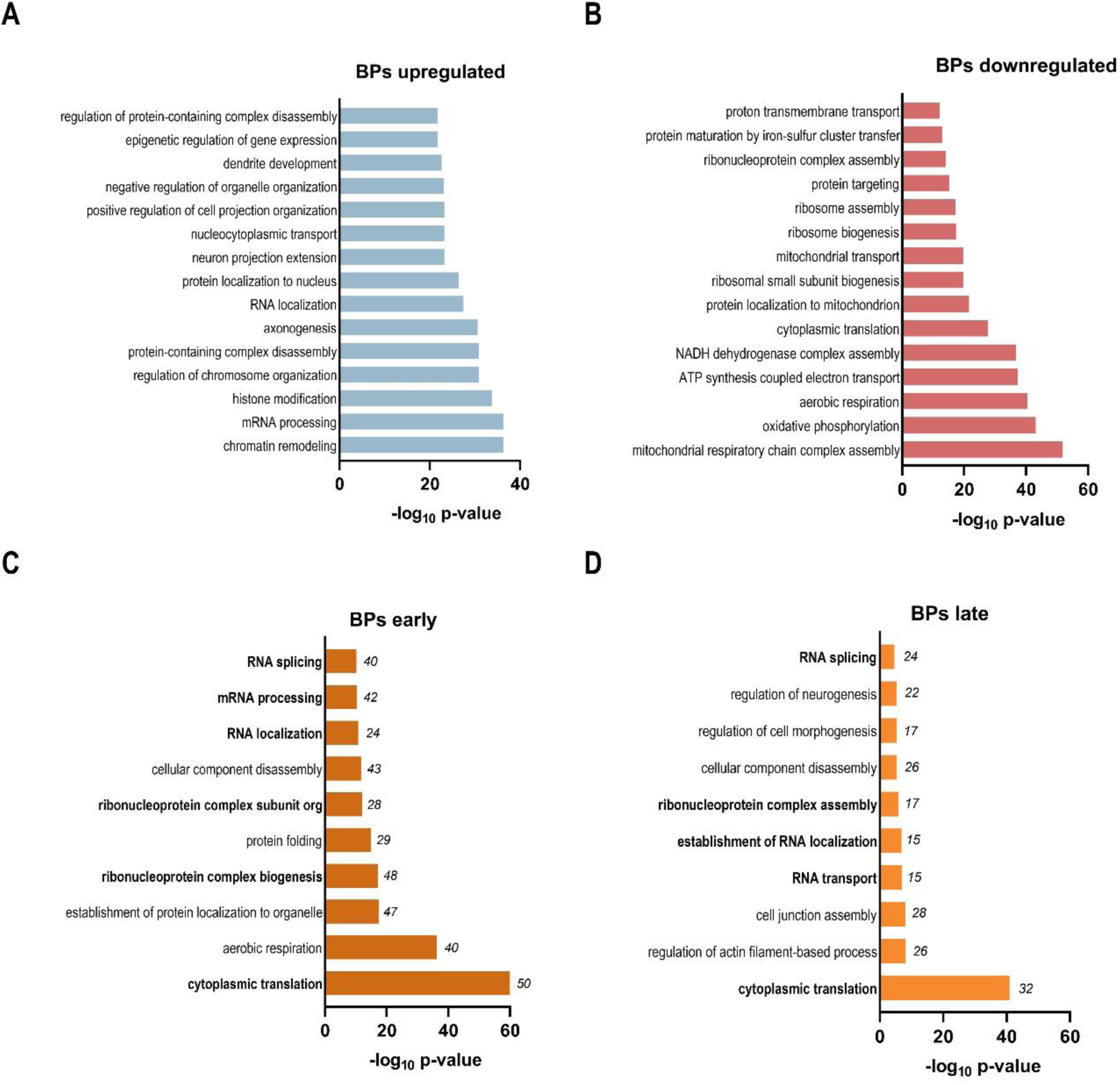
Analysis of functional transcriptional changes in oligodendrocytes in AD. (A, B) Gene ontology enrichment of Biological Process (BP) of upregulated genes (blue) and downregulated genes (red) *in vitro* after 24 hours of Aβ treatment. (C, D) Gene ontology enrichment analysis (BP only) shared in our Aβ-treated oligodendrocytes and early/late-AD conditions from Mathys et al (2019) (46).

**Table S2.** Characteristic of controls and AD subjects, categorised as stages I to VI of Braak and Braak and A, B or C of CERAD criteria.

| Case n<sup>o</sup> | Ref. n<sup>o</sup> | Braak stage<br>NFT | CERAD<br>Senile<br>plaque | Gender | Age | Region<br>analysed | Postmortem<br>delay* |
| --- | --- | --- | --- | --- | --- | --- | --- |
| 1 | 695 | I |  | M | 80 | Hp | 10:00 |
| 2 | 1378 | - |  | M | 78 | Hp | 6:00 |
| 3 | 1648 | - |  | M | 73 | Hp | 6:10 |
| 4 | 1357 | II |  | F | 79 | Hp | 10:30 |
| 5 | 1912 | II | B | F | 72 | Hp | 13:35 |
| 6 | 1937 | II | B | F | 83 | Hp | 7:20 |
| 7 | 1247 | III | A | F | 80 | Hp | 8:00 |
| 8 | 1345 | III | A | M | 78 | Hp | 8:00 |
| 9 | 1112 | III | B | F | 83 | Hp | 7:30 |
| 10 | 608 | IV | - | M | 78 | Hp | 7:00 |
| 11 | 1040 | IV | C | M | 76 | Hp | 8:25 |
| 12 | 1417 | IV | - | F | 79 | Hp | 4:30 |
| 13 | 977 | VI | C | M | 75 | Hp | 10:00 |
| 14 | 999 | VI | C | F | 76 | Hp | 10:00 |
| 15 | 1135 | VI | C | M | 79 | Hp | 6:25 |

**Figure S2.**
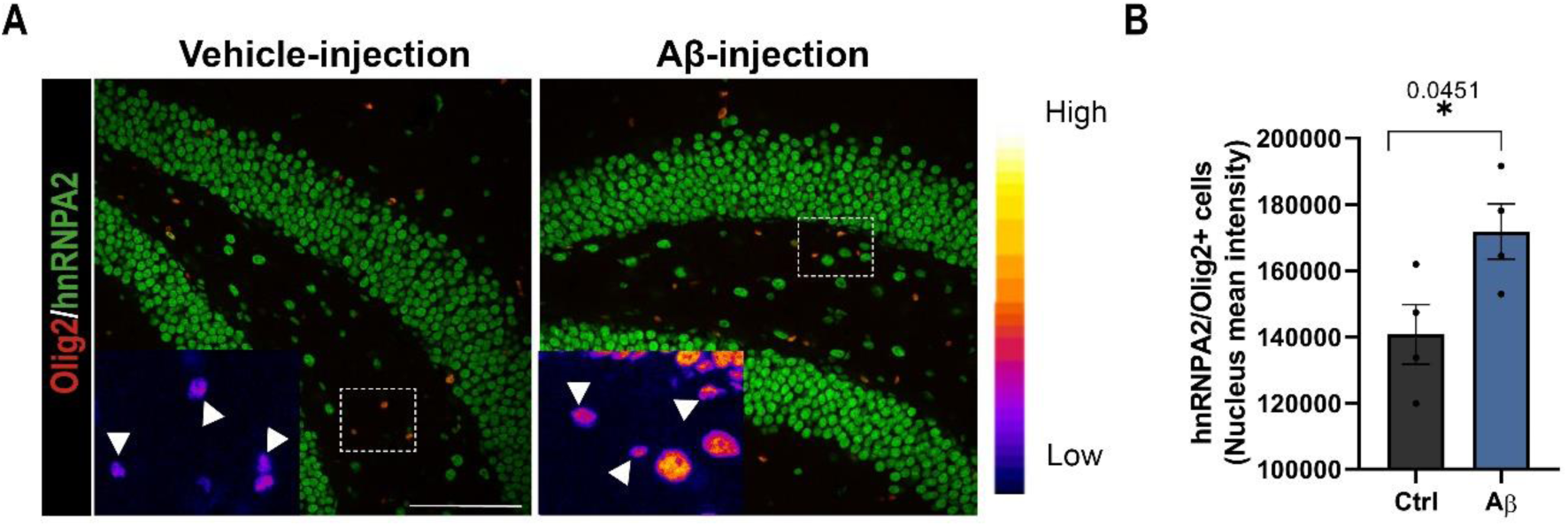
Analysis of hnRNP A2 expression in Aβ oligomer-injected mice. (A) Representative confocal images of Olig2 (red) and hnRNP A2 (green) in the dentate gyrus of vehicle and Aβ-injected mice. (B) Quantification of hnRNP A2 intensity in Olig2+ cells in the dentate gyrus. Scale bar 100 µm. Data indicate means ± S.E.M and dots represent individual animals, *p<0.05, compared to vehicle-injected mice. Statistical significance was drawn by two-tailed unpaired Student t-test.

**Figure S3.**
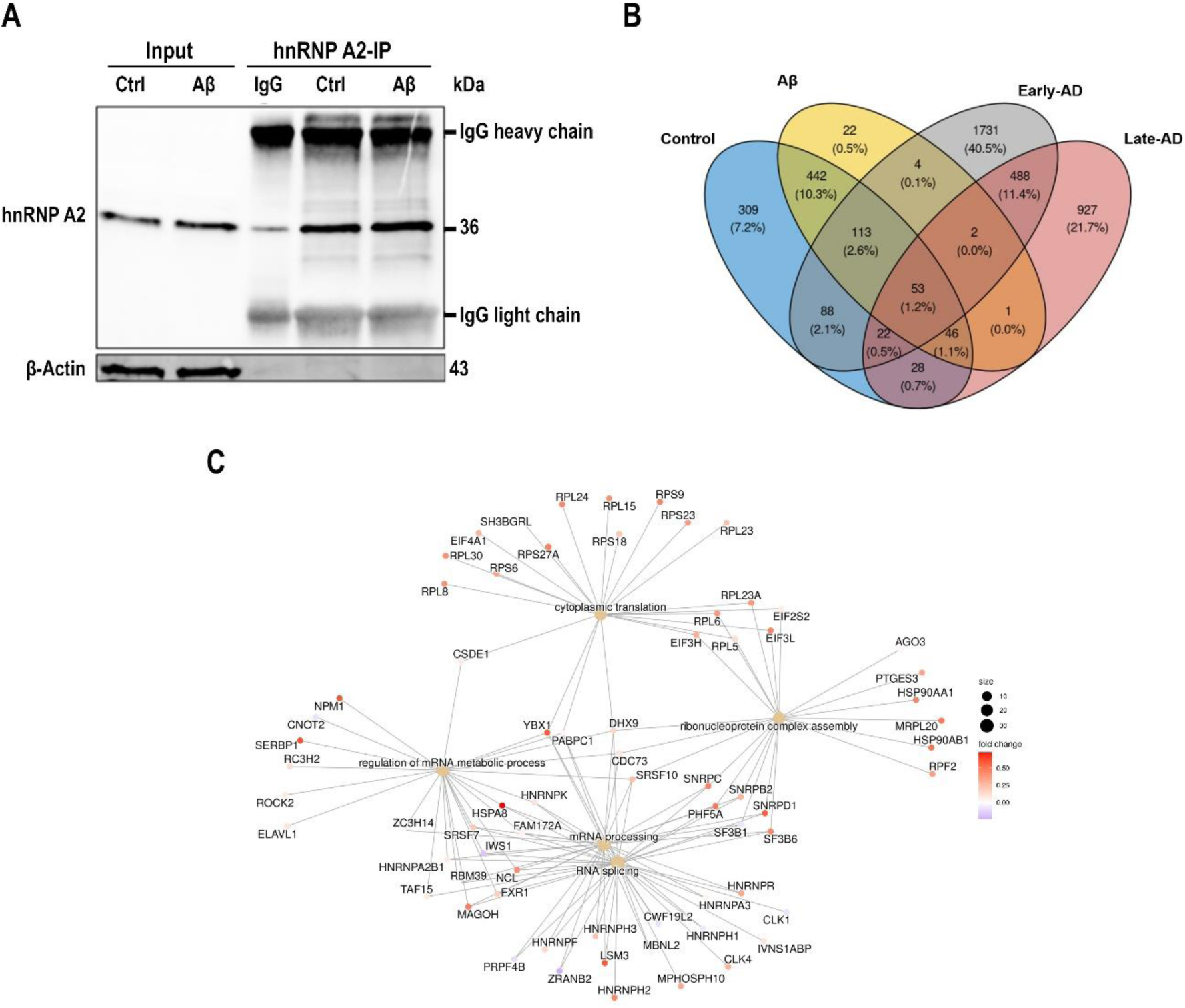
RIP-seq analysis of hnRNP A2 interactome performed in primary cultured oligodendrocytes compared to human gene signatures of AD. (A) IP was performed using hnRNP A2 antibody-coated agarose beads. Negative control of isotype IgG was used to detect non-specific binding of proteins to antibody. Proteins were eluted and loaded in 10% SDS-PAGE gel for analysis by western blot. (B) Venn diagram depicting the overlap between the hnRNP A2 interactome of control and Aβ-treated oligodendrocytes and the DEGs in human early and late-AD patients (46). Percentage and numbers indicate the genes shared among the conditions. (C) Top 5 GOs CNETPlot. Network visualization of DEGs involved in the top 5 enriched GO terms. Each gene is linked with its respective GO term or terms, if shared. The foldChange (from early-AD) is indicated colorwise and the amount of genes differentially expressed per term is shown sizewise for each node.

**Figure S4.**
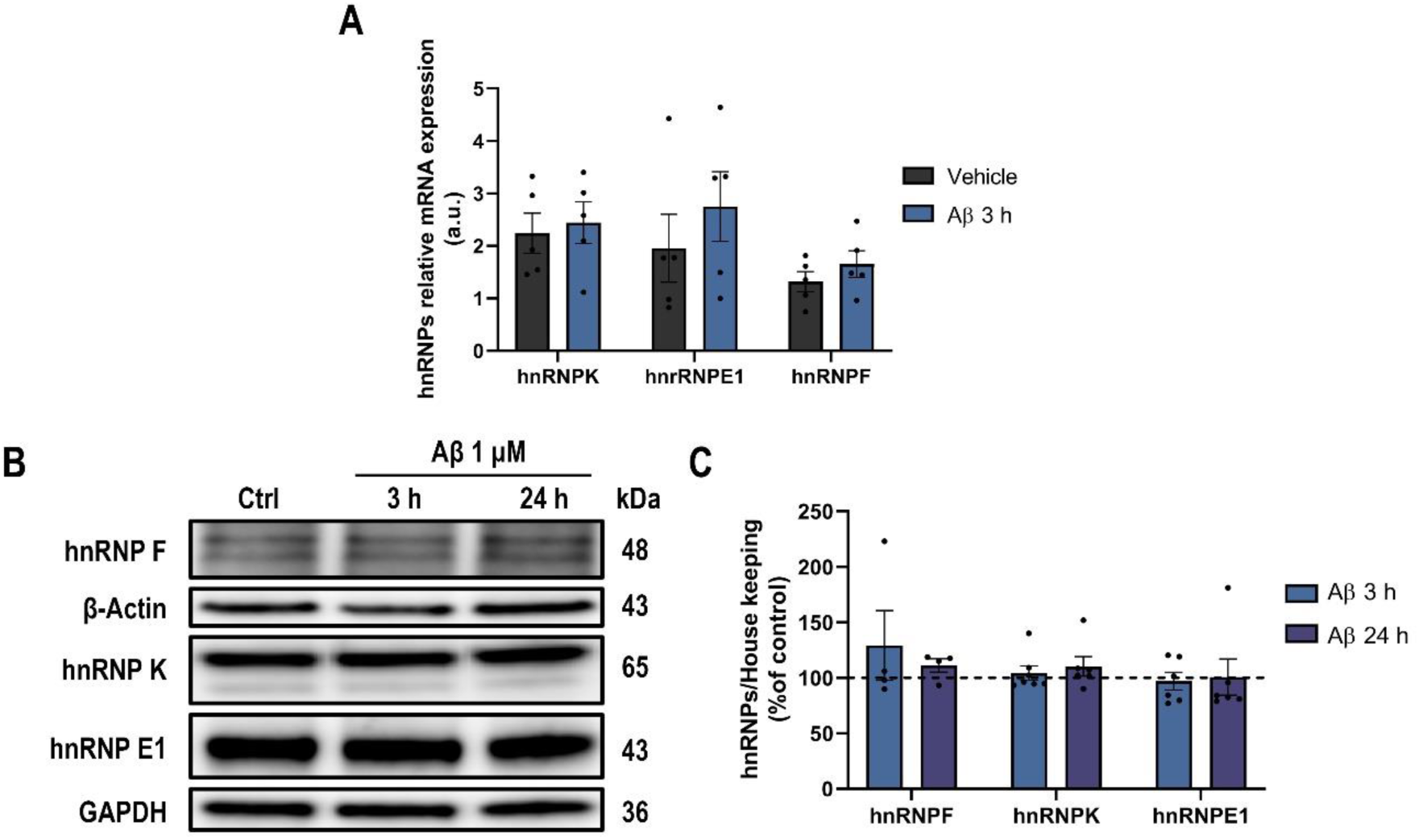
Analysis of hnRNPs found in *Mbp* and *Mobp* mRNA granule. (A) RT-qPCR analysis of hnRNPS in Aβ-treated and control cells. (B, C) hnRNPs western blot and relative quantification in total cell extracts from oligodendrocytes. Data indicate means ± S.E.M and dots represent independent experiments. Statistical significance was drawn by two-tailed paired Student’s t-test (A) and one-way ANOVA followed by Dunnett post-hoc test (C).

**Figure S5.**
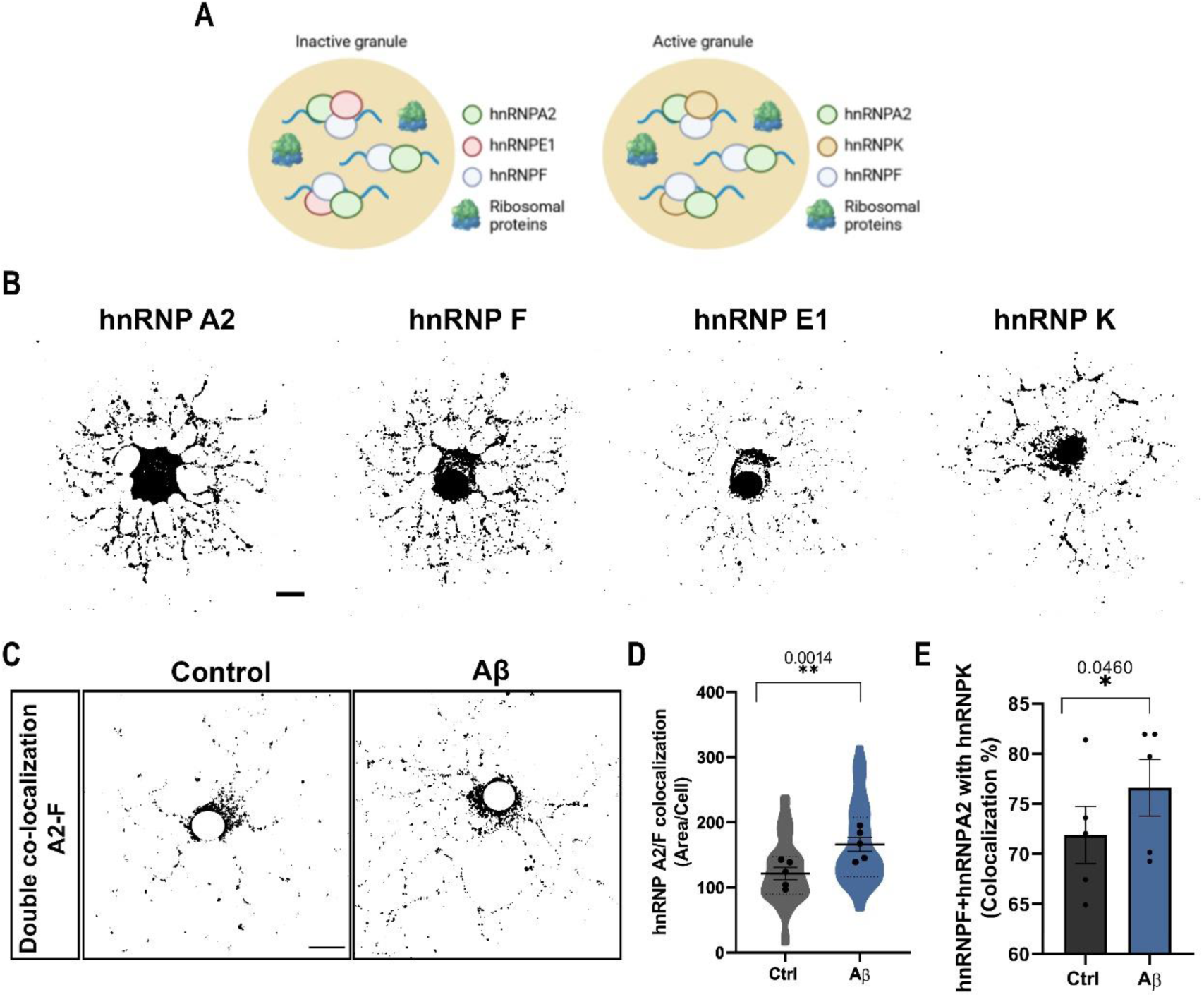
mRNA granule description and dynamic analysis. (A) During granule transport, MBP mRNA is maintained on a translationally silenced (inactive) or activated (active) states. (B) Representative micrographs of oligodendrocytes showing the intracellular localisation of the different hnRNPs. (C, D) Double colocalisation images of hnRNP A2 and hnRNP F. 1 µM Aβ significantly increased hnRNPA2/F colocalisation (µm^2^). Scale bar, 10 µm. (E) Graphs show that 70% of all granules contain hnRNPK (active granules) and Aβ-treated cells contain 5 % more active granules. (F, G) hnRNP A2 CO-IP with the different hnRNPs found in the granules. 1 µM Aβ exposure significantly increasedhnRNPK within the granule. Data indicate means ± S.E.M and dots represent independent experiments. Statistical significance (*p<0.05, **p<0.01) was drawn by two-tailed paired Student’s t-test (D, E).

**Figure S6.**
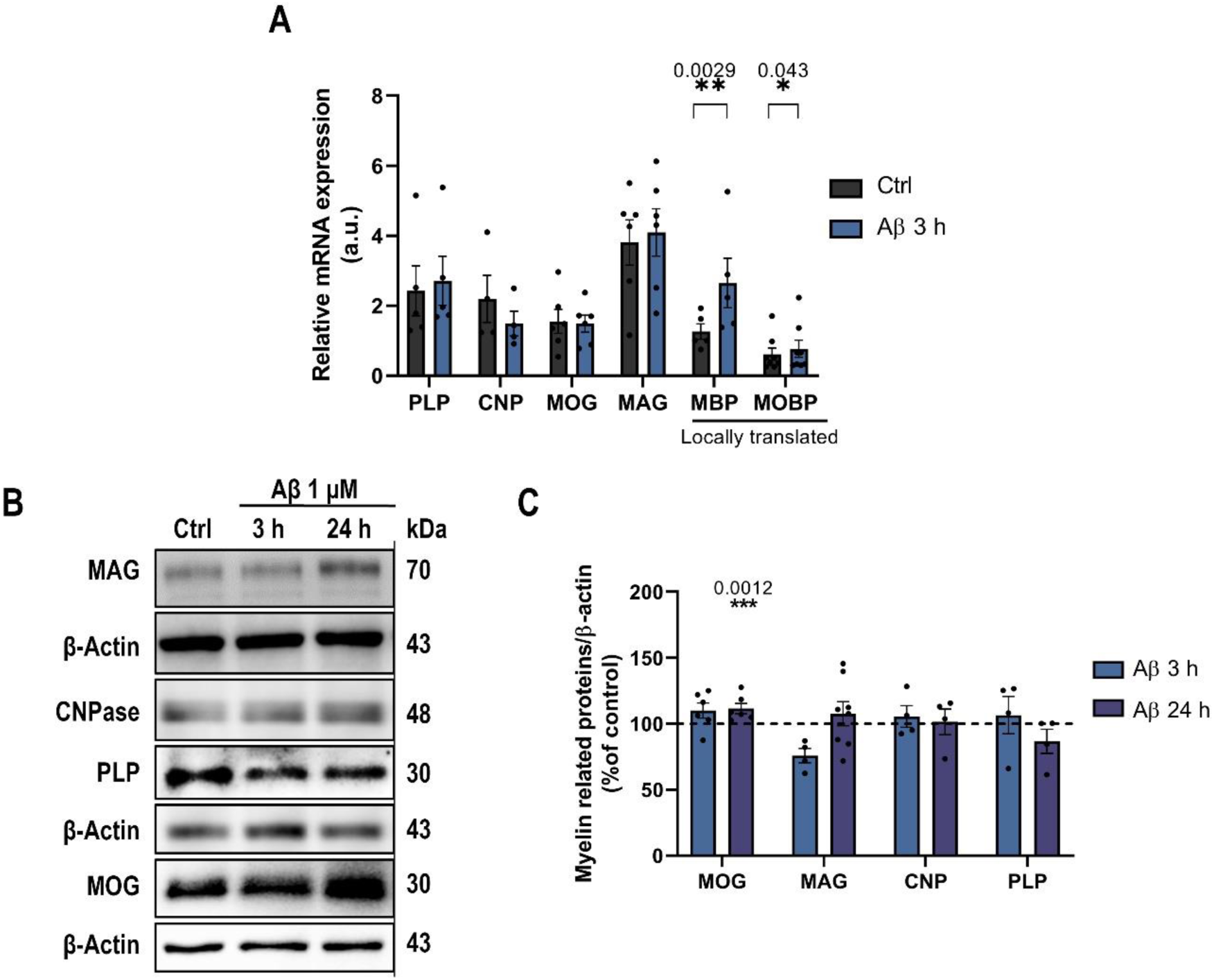
Aβ oligomer upregulate myelin-related proteins at mRNA and protein levels. (A) RT-qPCR analysis of different myelin-related proteins in treated and control cells. (B, C) MBP and MOBP expression and relative quantification in treated and control oligodendrocyte cell extracts. (D, E) Different myelin related protein expression and relative quantification in treated and control oligodendrocyte cell extracts. Data are represented as means ± S.E.M and dots indicate independent experiments, *p<0.05, **p<0.01, ***p<0.001 compared to control cells. Statistical significance was drawn by two-tailed paired Student’s t-test (B) and one-way ANOVA followed by Dunnett post-hoc test (C).

**Figure S7.**
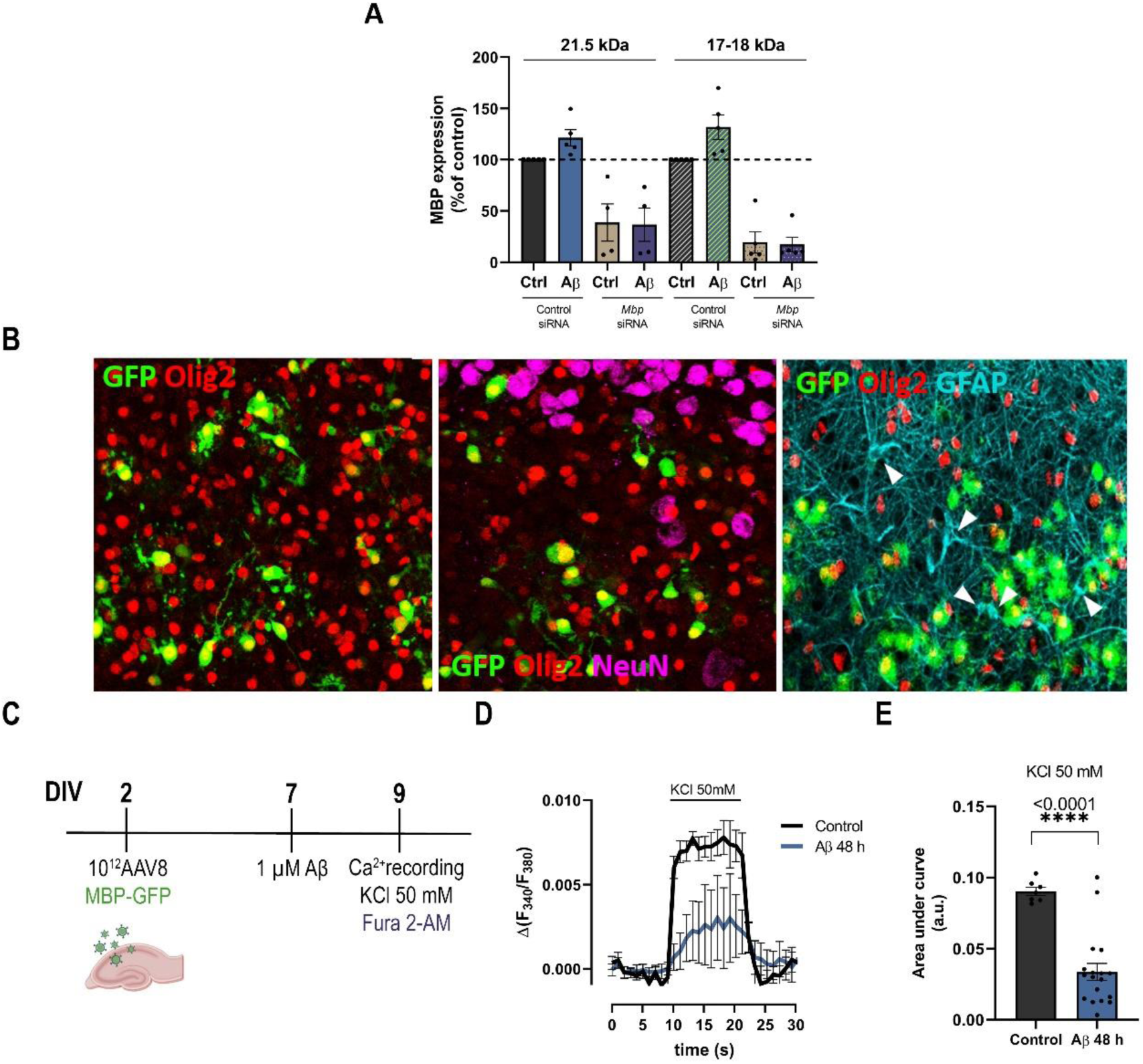
Aβ oligomers inhibit intracellular calcium influx in oligodendrocytes of hippocampal organotypic slices. (A) Western blot and analysis of MBP expression levels following Aβ exposure in oligodendrocytes treated with control siRNA and *Mbp* siRNA are shown. (B) Representative confocal z-stacks projections of hippocampal organotypic slices infected with AAV8-MBP-GFP (green) immunostained with Olig2 (red), NeuN (magenta) and GFAP (cyan). Note that only olig2 colocalized with GFP. (B) Scheme of the experimental approach. (C) Cells were loaded with Fura 2-AM and KCl-induced intracellular calcium levels were recorded. (D) Histogram depicting area under the curve (AUC) obtained by KCl 50 mM application in vehicle-and Aβ-treated hippocampal organotypic slices. Data indicate means ± S.E.M and dots represent individual cells, ****p<0.0001, compared to vehicle-treated organotypic oligodendrocytes. Statistical significance was drawn by two-tailed unpaired Student’s t-test.

